# Human pluripotent stem cell-derived intestinal epithelial cells maintain small intestine-specific functions over time, even with repeated cell division

**DOI:** 10.1101/2025.02.17.638577

**Authors:** Junlong Chen, So Kuramochi, Shinichiro Horiuchi, Tomoyuki Kawasaki, Hiroto Kakizaki, Lirika Tabata, Tohru Kimura, Kazuaki Nakamura, Hidenori Akutsu, Seiichi Ishida, Atsuo Kikuchi, Akihiro Umezawa

**Author notes:** These two authors contributed equally to this work (Junlong Chen, So Kuramochi).

## Abstract

The human colon cancer-derived cell line Caco-2 is widely used in drug discovery due to its barrier function and transporter activity. However, Caco-2 cells have extremely low drug metabolic capacity, resulting in discrepancies with human physiology. In this study, we conducted experiments on human intestinal epithelial cells generated from pluripotent stem cell-derived organoids. We assessed cell morphology, gene expression, barrier and transporter functions, drug metabolic capacity, and cytotoxicity in relation to cell growth and the effects of cellular aging. The results indicate that organoid-derived intestinal epithelial cells may be helpful as a new model cell for drug discovery. Understanding the advantages of drug metabolic capacity and cytotoxicity among cryopreserved human enterocytes, the human colon cancer-derived cell line Caco-2, and human pluripotent stem cell-derived intestinal epithelial cells within microphysiological systems and organ-on-chip technologies is essential for the development of an appropriate model system for the small intestine.

## 1. Introduction

The small intestine is the largest organ in the human body and is responsible for absorbing nutrients and transporting them through the bloodstream to the entire body. Intestinal epithelial cells are one of the functional units of the small intestine, handling the absorption of nutrients and water while maintaining intestinal mechanisms that prevent foreign substances from entering the body. These intestinal functions include barrier function, transporter activity, and drug metabolic capacity^1,2^. Since the body considers pharmaceuticals foreign substances, these intestinal functions limit their entry into the body. For pharmaceuticals to exhibit their therapeutic effects, absorption from intestinal epithelial cells is indispensable, making small intestine in vitro models highly sought after for drug discovery screening, pharmacokinetic studies, bioavailability prediction, and toxicity and safety evaluations^3–5^. The human colon cancer-derived cell line Caco-2 is frequently used as a small intestine in vitro model. While Caco-2 cells possess barrier function and transporter activity, their drug metabolic capacity is extremely low. In particular, they are deficient in CYP3A4, which is involved in the metabolism of many pharmaceuticals, and CES2, which is involved in the metabolism of prodrugs, leading to discrepancies with the human small intestine^6,7^. Therefore, developing new small intestine in vitro models, in addition to Caco-2 cells, is necessary.

One in vitro model of the small intestine is the three-dimensional intestinal organoid that mimics the structure of the small intestinal mucosa. Intestinal organoids are derived from crypts, have a single-layer structure, and contain differentiated cells such as stem cells, transit-amplifying (TA) cells, absorptive epithelial cells, and goblet cells^8^. Organoid culture enabled the proliferation of intestinal epithelial cells in vitro for the first time. Additionally, differentiation from human pluripotent stem cells to intestinal organoids, mimicking the process of embryonic development, has been reported^9,10^. However, the lumen of intestinal organoids faces inward, making them unsuitable for drug administration experiments. Therefore, the 2D culture of intestinal epithelial cells is necessary for drug discovery research. Differentiation from human pluripotent stem cells to intestinal epithelial cells in 2D culture has been reported^11–13^. Moreover, there have been reports of transferring three-dimensional intestinal organoids to adherent culture^14^. However, in these models, intestinal epithelial cells, after adherent culture, do not possess proliferative capacity, and due to the long differentiation period, it is challenging to supply cells rapidly and in large quantities.

Furthermore, there is the issue of functional lot-to-lot variability. We reported that intestinal epithelial cells can be proliferatively cultured in 2D using mesenchymal stromal cells^15^. Intestinal epithelial cells and mesenchymal cells are derived from intestinal organoids, referred to as organoid-derived intestinal epithelial cells and organoid-derived mesenchymal stromal cells, respectively. Under the support of organoid-derived mesenchymal stromal cells, organoid-derived intestinal epithelial cells proliferate in 2D and can be subcultured and cryopreserved. Furthermore, organoid-derived intestinal epithelial cells maintain intestinal functions during proliferation, exhibit morphologically columnar epithelial cells, and express markers of small intestinal epithelial cells. Additionally, they possess barrier function, transporter activity, and drug metabolic capacity, which are critical intestinal functions in drug discovery research.

This study investigated the impact of cellular aging on the function of organoid-derived intestinal epithelial cells to evaluate their suitability for drug discovery, particularly in assessing the effects of long-term drug administration for safety evaluations. We developed a system to culture model cells long-term while maintaining functionality, demonstrating their potential as standard drug discovery tools by evaluating barrier function, transporter activity, and drug metabolism.

## 1. Materials and methods

### Culture of human iPSCs

Human iPSCs (EDOM22 #8) derived from menstrual blood were used^16,17^. The iPSCs were cultured on dishes coated with vitronectin (A14700, Thermo Fisher Scientific) in Stem flex medium (A3349401, Thermo Fisher Scientific). The dishes were coated with 5 μg/ml vitronectin at room temperature for 1 h, and the medium was changed daily. The iPSCs were detached with 0.5 mM EDTA (06894–14, Nacalai tesque) for 5 min and passaged at 1100 cells/cm^2^ weekly.

### Culture of intestinal organoid-derived mesenchymal stromal cells

Intestinal organoid-derived mesenchymal stromal cells derived from Mini-Guts were used^15,18^. The intestinal organoid-derived mesenchymal stromal cells were cultured on non-coating dishes at 37°C in 5% CO_2_ in DMEM (D6429, Sigma-Aldrich) supplemented with 10% fetal bovine serum (FBS) (10,091,148, Thermo Fisher Scientific) to allow cell growth. The medium was changed weekly. The intestinal organoid-derived mesenchymal stromal cells were detached with 0.25% trypsin/1 mM EDTA (209–16941, Fujifilm Wako Pure Chemicals Co., Ltd.) for 3 min and passaged into new dishes or cryopreserved with STEM-CELLBANKER (CB045, Nippon Zenyaku Kogyo Co. Ltd.) until use.

### Culture of organoid-derived intestinal epithelial cells

Organoid-derived intestinal epithelial cells derived from Mini-Guts were used ^15,18^. The organoid-derived intestinal epithelial cells were cultured on dishes with intestinal organoid-derived mesenchymal stromal cells at 37°C in 5% CO2 in EMUKK-05 medium (Mx) to allow cell growth. The dishes were confluent with intestinal organoid-derived mesenchymal stromal cells. The medium was changed every 2 days. The organoid-derived intestinal epithelial cells were detached with 0.25% trypsin/1 mM EDTA for 5 min and passaged into 4 dishes or cryopreserved with STEM-CELLBANKER until use.

### Caco-2 Cell Culture

Caco-2 cells were purchased from ECACC (Lot: 12F018). They were cultured in Dulbecco’s Modified Eagle Medium (DMEM) supplemented with 10% FBS at 37°C in 5% CO2 for 14 days, with the medium changed every 2 days. The cells were seeded at 2.0 × 105 cells/cm2.

### Karyotypic analysis

Karyotypic analysis was contracted to Nihon Gene Research Laboratories (Sendai, Japan). To assess diploidy, 50 cells at metaphase were examined. Metaphase spreads were prepared from cells treated with 100 ng/mL of Colcemid (KaryoMax, Gibco; Thermo Fisher Scientific, MA, USA) for 6 h. The cells were fixed with methanol: glacial acetic acid (2:5) three times and placed onto glass slides. Giemsa banding was applied to metaphase chromosomes. A minimum of 10 metaphase spreads were analyzed for each sample and karyotyped using a chromosome imaging analyzer system (Applied Spectral Imaging, CA, USA).

### Cell proliferation analysis

The intestinal epithelial cell proliferation was analyzed using time-lapse photography and WST-1 Cell Proliferation Assay System (MK400, TAKARA BIO INC.). The time-lapse filming started on the day of the epithelial cell seeding using the Incucyte S3 (SARTORIUS). Analyses using the WST-1 Cell Proliferation Assay System were started the day after organoid-derived intestinal epithelial cells seeding and were analyzed twice a week until day 48. The absorbance of the organoid-derived intestinal epithelial cells was determined by subtracting the absorbance of intestinal organoid-derived mesenchymal stromal cells only.

### qRT-PCR

The cultured cells were washed twice with Dulbecco’s phosphate-buffered saline (Sigma-Aldrich, St Louis, MO, USA) before the total RNA was isolated. RNA extraction was performed using the RNeasy total RNA extraction kit (QIAGEN, Hilden, Germany) per the manufacturer’s guidelines. To quantify the gene expression levels, qPCR was performed using 8 ng of total RNA, following reverse transcription with the High Capacity RNA-to-cDNA kit (Thermo Fisher Scientific, Waltham, MA, USA) according to the manufacturer’s protocol. The QuantStudio 7 Flex Real-Time PCR system (Applied Biosystems, Foster City, CA, USA) was used to measure gene expression levels, and primers and probe sets were utilized to detect each gene transcript, as listed in Table 1. The expression levels of the genes are presented relative to RNAs derived from a human small intestine (R1234226-50, BioChain Institute, Inc., Newark, CA, USA).

**Table 1.** Primers used for qRT-PCR.

| Gene Symbol | Assay ID |
| --- | --- |
| CYP1A1 | Hs01054797_g1 |
| CYP2B6 | Hs00167927_g1 |
| CYP2C9 | Hs00426397_m1 |
| CYP2C19 | Hs00426380_m1 |
| CYP2D6 | Hs00164385_m1 |
| CYP3A4 | Hs00430021_m1 |
| P-gp | Hs00184500_m1 |
| BCRP | Hs01053790_m1 |
| PEPT1 | Hs00192639_m1 |
| CES1 | Hs00275607_m1 |
| CES2 | Hs01077945_m1 |

Assay ID is the name of the probe and primer of the TaqMan Assay.

### Preparation for histological studies

Samples were coagulated in iPGell (PG20-1, GenoStaff) following the manufacturer’s instructions and fixed in 4% paraformaldehyde at 4°C overnight. Fixed samples were embedded in a paraffin block to prepare thin cell sections. Deparaffinization, dehydration, and permeabilization were performed using standard techniques. Hematoxylin-eosin (HE) staining was performed with Carrazzi’s hematoxylin solution (30,022, Muto Chemicals) and eosin Y (32053, Muto Chemicals). Alcian blue staining was achieved with alcian blue solution pH2.5 (40,852, Muto Chemicals). Nuclei were counterstained with kernechtrot solution (40872, Muto chemicals).

### Immunostaining

Cells in culture (3910-035, IWAKI) were fixed with 4% paraformaldehyde for 15 min at room temperature. After washing with phosphate-buffered saline (PBS), cells were permeabilized with 0.25% Triton X-100 in PBS for 20 min, pre-incubated with Protein Block Serum-Free (X0909, Dako) for 30 min at room temperature, and then exposed to primary antibodies overnight at 4°C. After three washes with PBS, the cells were incubated with fluorescent secondary antibodies for 1 h at room temperature and washed three times with PBS. Nuclei were stained with a mounting medium containing 4’,6-diamidino-2-phenylindole dihydrochloride solution (DAPI) (H-1200, Vector Laboratories).

**Table 2.**
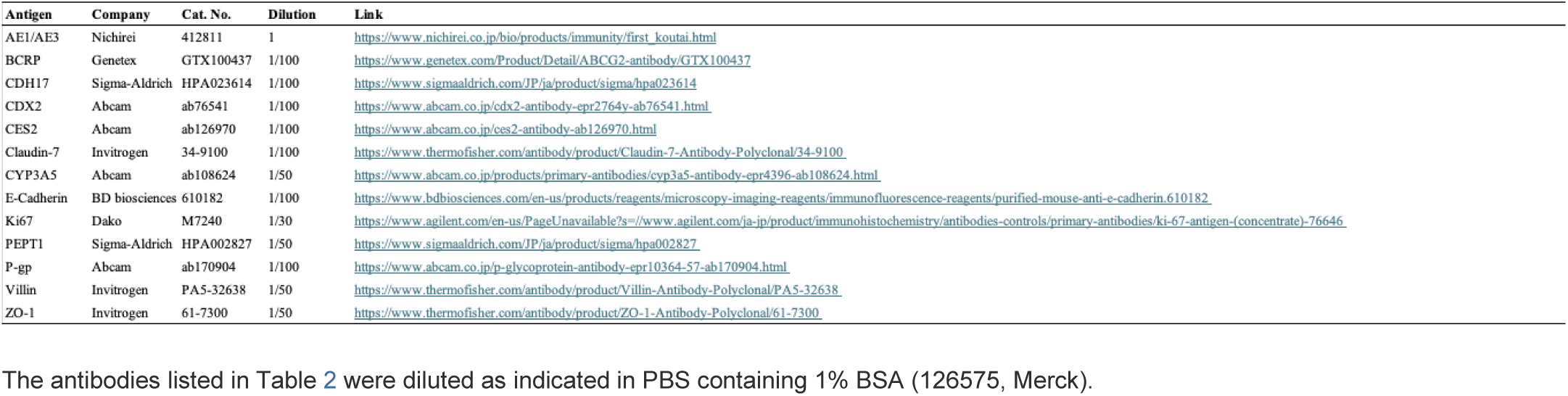
Antibodies used for immunostaining.

**Table 3.**
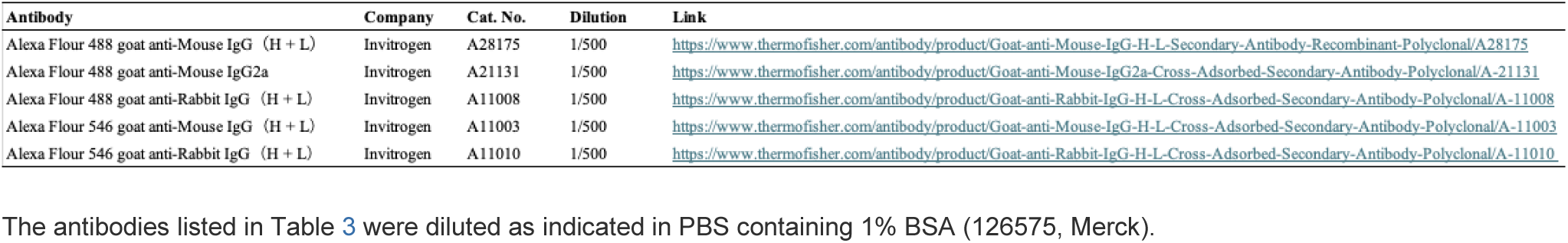
Secondary antibodies used for immunostaining.

### Transmission electron microscopy (TEM) and scanning electron microscopy (SEM)

Cultured cells were washed three times with PBS for transmission electron microscopy. Fixation was performed in PBS containing 2.5% glutaraldehyde for 2 h. The cells were embedded in epoxy resin. Ultrathin sections cut vertically to the culture surface were double stained in uranyl acetate and lead citrate and were viewed under a JEM-1200 PLUS transmission electron microscope (Nihon Denshi) and SU6600 (Hitachi High-Tech corporation).

### Trans-epithelial electrical resistance (TEER) measurements

The cells were seeded onto Transwell inserts (353095, Falcon) and cultured until confluent. The Millicell-ERS (Electrical Resistance System) with chopstick electrodes was used to measure construct trans-epithelial electrical resistance (TEER). The instrument was calibrated with a 1000 Ω resistor before measurements, and an empty Transwell was used as a blank. Constructs were equilibrated to room temperature and switched to basal media for readings. The blank Transwell and all samples were measured three times. Samples and blanks were measured in triplicate, averaged, and used in the following calculations:

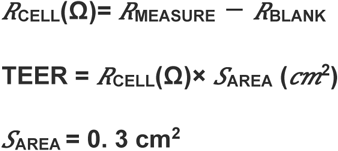

### Permeability assay

The cells were seeded on Transwell inserts and cultured until confluent. Before the start of the assay, samples were pre-incubated with transport buffer (Hank’s Balanced Salt Solution (HBSS) (14025092, Gibco) supplemented with 10 nM HEPES (15630106, Gibco) and 4.5 mg/ml glucose) for 1 h. Lucifer yellow solution (125-06281, Fujifilm Wako Pure Chemicals Co. Ltd.) was added to the apical side to a final concentration of 300 μM and then drawn out of the basal side at 30 min intervals for up to 120 min at 37°C with shaking at 40 rpm. Lucifer yellow fluorescence was measured using a SYNERGY H1 microplate reader (Bio-Tek) at 428 nm excitation and 536 nm emission. Papp was calculated using the following equation:

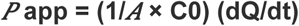

where dQ/dt is the drug permeation rate across the cell monolayer, C0 is the donor-compartment concentration at time zero, and A is the area of the cell monolayer.

### Transport assay

The cells were seeded on Transwell inserts and cultured until confluent. Before the start of the assay, samples were pre-incubated with transport buffer for 1 h. Rhodamine123 solution (187-01703, Fujifilm Wako Pure Chemicals Co. Ltd.) was added to the apical or basal side to a final concentration of 5 μM and then drawn out of the basal side at 60 min intervals for up to 120 min at 37°C with shaking 40 rpm. After each sampling, an equal volume of transport buffer was immediately added to the drawn-out side. Zosuqidar solution (21533, Fujifilm Wako Pure Chemicals Co., Ltd.) was added as an inhibitor at a final concentration of 5 μM. FL Prazosin solution (B7433, Thermo Fisher Scientific) was added to the apical or basal side to a final concentration of 10 μM and then drawn out of the basal side at 60 min intervals for up to 120 min at 37℃ with shaking at 40 rpm. After each sampling, an equal volume of transport buffer was immediately added to the drawn-out side. Ko143 solution (K2144, Sigma-Aldrich) was added as an inhibitor at a final concentration of 10 μM. Inhibition using the Ko 143 solution and Zosuquidar solution was also performed. Rhodamine123 fluorescence was measured using a SYNERGY H1 microplate reader at 480 nm excitation and 530 nm emission, and Prazosin fluorescence was measured at 490 nm excitation and 520 nm emission.

The ER was defined as Papp (basal-to-apical)/Papp (apical-to-basal).

### Cytochrome P450 induction assays

Organoid-derived intestinal epithelial cells cultured in the 24-well plates in EMUKK-05 medium were treated with 100 nM vitaminD3 (D1530, Sigma-Aldrich) or 50 μM omeprazole (150-02091, Fujifilm Wako Pure Chemicals Co. Ltd.) for 24 h or 20 μM rifampicin (189-01001, Fujifilm Wako Pure Chemicals Co. Ltd.) or 500 μM phenobarbital (162-11602, Fujifilm Wako Pure Chemicals Co. Ltd.) or 100 μM dexamethasone (194561, MP Biomedicals) for 48 h at a cell density of 80%. Vehicles were treated with DMSO (043-07216, Wako Pure Chemical Industries, Ltd., Japan) (final concentration 0.1%).

### Determination of cytochrome P450 activity

The cells were incubated with the assay buffer (HBSS supplemented with 10 nM HEPES and 4.5 mg/ml glucose) containing 500 μM bupropion (028-17311, Fujifilm Wako Pure Chemicals Co., Ltd.), 25 μM diclofenac (043-22851, Fujifilm Wako Pure Chemicals Co., Ltd.), 250 μM (S)-Mephenytoin (518-20371, Fujifilm Wako Pure Chemicals Co., Ltd.), 100 μM Bufuralol (UC168, Sigma-Aldrich), and 20 μM Midazolam (135-13791, Fujifilm Wako Pure Chemicals Co., Ltd.) at 37°C for 2 h and the assay medium was withdrawn at 30 and 120 min. The cells were finally detached with 0.25% trypsin/1 mM EDTA and collected for cell counting. The amount of these metabolites, hydroxybupropion, 4′-Hydroxydiclofenac, 4′-Hydroxymephenytoin, 1′-Hydroxybufuralol, and 1′-Hydroxymidazolam were measured by LC-MS/MS (InertSustain® AQ-C18, LC-20A (SHIMADZU), API4000 (AB Sciex Pte. Ltd.)). Sumika Chemical Analysis Service, Ltd performed the measurements.

### Toxicity assays (Naproxen and Acetaminophen)

The cells were incubated with the culture medium containing Naproxen (147-07201, Fujifilm Wako Pure Chemicals Co. Ltd.) or Acetaminophen (A7085, Sigma-Aldrich) and 1% DMSO at 37°C for 1 day. The WST-1 Cell Proliferation Assay System finally performed the cells for cell viability counting. The absorbance of the untreated cells was set as 100% viability, and the cell viability of the vehicle and test conditions were determined.

### RNAseq analysis

All statistical analyses were performed using R version 4.3.3-4.4.0. Packages used included edgeR, dplyr, ggplot2, genefilter, clusterProfiler for Gene Ontology (GO) analysis, org.Hs.eg.db, and ComplexHeatmap. Read count data was used for differential gene expression (DEG) and Gene Ontology analyses. Lowly expressed genes were filtered using the filterByExpr() function from the edgeR package. Normalization factors were calculated using calcNormFactors(), estimateCommonDisp(), and estimateTagwiseDisp() functions in edgeR. Normalized expression values were then output as Counts Per Million (CPM) using the cpm() function from edgeR. A correlation heatmap was generated using all genes to clarify the differences between passages. Principal Component Analysis (PCA), dendrogram, and correlation plot were generated using a subset of genes associated with intestinal functions, selected based on a gene list (table4), to clarify the changes in intestinal functions. Two-sample comparison tests were performed using the exactTest() function from the edgeR package, and volcano plots were generated to visualize differential gene expression between Passage 6 vs. Passage 9 and Passage 6 vs. Passage 12, displaying log2-fold changes and False Discovery Rate (FDR) values. Highly variable genes were selected, multiple comparison test was performed to 3 groups and genes with an FDR less than 0.05 were extracted, and GO enrichment analysis was performed using the enrichGO() function from the clusterProfiler package. Redundant GO terms were simplified using the simplify() function from clusterProfiler, and barplots were created using the barplot function from the enrichplot package to visualize enriched GO terms. Two-group comparison tests were performed for three pairwise comparisons: P6 vs. other, P9 vs. other, and P12 vs. other. A heatmap was created using genes with an FDR < 0.05 and a fold change greater than 3.0 in respective groups. Multiple comparison tests were performed across the three groups (P6 vs. P9 vs. P12). A Likelihood Ratio Test was performed using the glmFit() and glmLRT() functions from the edgeR package. The Benjamini-Hochberg method was applied to correct for multiple comparisons, and the FDR was obtained. A heatmap was created using genes with an FDR < 0.05 and a fold change greater than 3.0 in respective groups. Finally, a heatmap of transporter genes was generated using the ComplexHeatmap package to visualize changes across passages.

## 2. Results

### Organoid–derived intestinal epithelial cells have a high proliferative capability

Intestinal epithelial cells were generated from organoids derived from human induced pluripotent stem cells (iPSCs). We evaluated the proliferative limits at which organoid-derived intestinal epithelial cells can maintain their cellular functions and assessed the impact of senescence. Since a decrease in cell proliferation was observed at Passage 12, we assessed the proliferation capacity, morphology, barrier function, transporter activity, drug metabolic capacity, and comprehensive gene expression of organoid-derived intestinal epithelial cells at Passage 6, 9, and 12. Organoid-derived intestinal epithelial cells exhibited similar microscopic images across all passages, with the cells being small and highly densely packed by observation under phase-contrast microscopy (Fig. 1A and S1).

**Figure 1.**
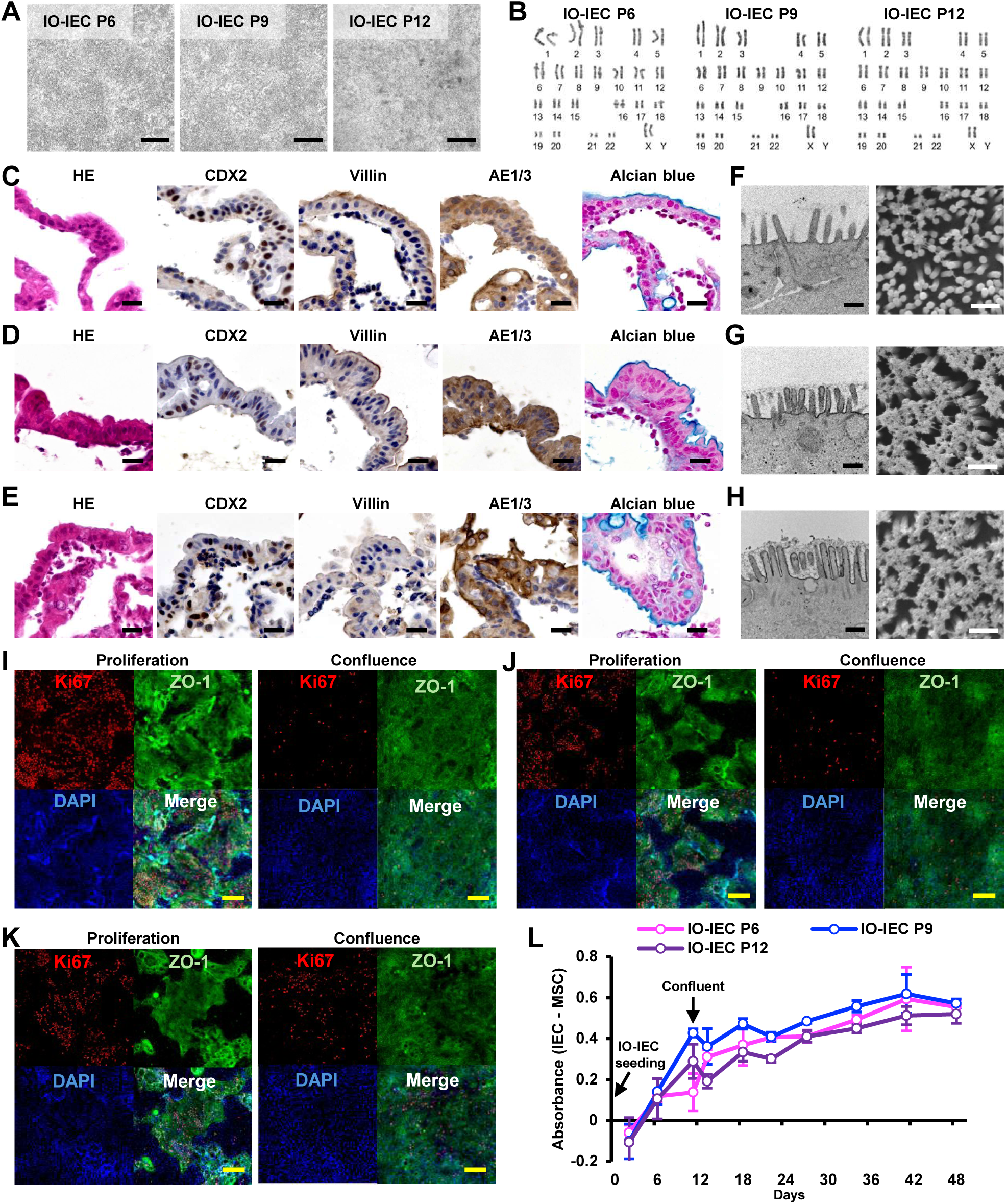
Proliferative potential and morphological analysis of organoid-derived intestinal epithelial cells. Intestinal organoid-derived intestinal epithelial cells are abbreviated as IO-IEC. A. Phase-contrast micrograph of organoid-derived intestinal epithelial cells at each passage. Scale bars: 500 µm. B. Karyotypic analysis revealed that organoid-derived intestinal epithelial cells at each passage have normal karyotypes. C. Histology and immunohistochemistry of intestinal epithelial cells (organoid-derived intestinal epithelial cells) at Passage 6. Immunohistochemistry was performed with antibodies to CDX2, Villin, and AE1/AE3. Scale bars: 20 µm. D. Histology and immunohistochemistry of organoid-derived intestinal epithelial cells at Passage 9. Immunohistochemistry was performed with antibodies to CDX2, Villin, and AE1/AE3. Scale bars: 20 µm. E. Histology and immunohistochemistry of organoid-derived intestinal epithelial cells at Passage 12. Immunohistochemistry was performed with antibodies to CDX2, Villin, and AE1/AE3. Scale bars: 20 µm. F. Transmission electron microscopy and scanning electron microscopic analysis of the apical membrane side of organoid-derived intestinal epithelial cells at Passage 6. Scale bars: 500 nm. G. Transmission electron microscopy and scanning electron microscopic analysis of the apical membrane side of organoid-derived intestinal epithelial cells at Passage 9. Scale bars: 500 nm. H. Transmission electron microscopy and scanning electron microscopic analysis of the apical membrane side of organoid-derived intestinal epithelial cells at Passage 12. Scale bars: 500 nm. I. Immunohistochemistry was performed with antibodies to Ki67 (proliferation marker) and ZO-1 (tight junction protein) in passage 6 organoid-derived intestinal epithelial cells. Cells were examined in proliferating and confluent conditions. Nuclei were counterstained with DAPI. Scale bars: 200 µm. J. Immunohistochemistry was performed with antibodies to Ki67 (proliferation marker) and ZO-1 (tight junction protein) in passage 9 organoid-derived intestinal epithelial cells. Cells were examined in proliferating and confluent conditions. Nuclei were counterstained with DAPI. Scale bars: 200 µm. K. Immunohistochemistry was performed with antibodies to Ki67 (proliferation marker) and ZO-1 (tight junction protein) in passage 12 organoid-derived intestinal epithelial cells. Cells were examined in proliferating and confluent conditions. Nuclei were counterstained with DAPI. Scale bars: 200 µm. L. Measurement of the proliferative capacity of organoid-derived intestinal epithelial cells using the WST-1 Cell Proliferation Assay System. The feeder cells’ intestinal organoid-derived mesenchymal stromal cells were subtracted to determine the absorbance value of organoid-derived intestinal epithelial cells alone.

Karyotypic analysis showed that these cells had chromosomal stability without any chromosomal abnormalities (Fig. 1B). Histological analysis indicated that organoid-derived intestinal epithelial cells exhibited typical columnar epithelial morphology at all passages and were positive for the small intestinal epithelial cell markers CDX2 and AE1/3 (Fig. 1C–E). Additionally, in the apical membrane of organoid-derived intestinal epithelial cells, the localization of Villin and the formation of microvilli and glycocalyx were observed, indicating that organoid-derived intestinal epithelial cells retained cellular polarity even after multiple passages (Fig. 1F–H).

With increasing passages, the length of microvilli and the density of the glycocalyx in organoid-derived intestinal epithelial cells increased, suggesting cellular maturation. In colonies of proliferating organoid-derived intestinal epithelial cells, most of the cells were Ki67-positive (Fig. 1I–K). Notably, the proportion of Ki67-positive cells before and after confluency decreased from 52.3% to 7.6% at Passage 6, from 60.5% to 5.2% at Passage 9, and from 27.0% to 19.0% at Passage 12 (Fig. S2).

Despite the reduction in Ki67-positive cells, cell proliferation continued without the disappearance of Ki67-positive cells, and the overall metabolic activity of the cells gradually increased (Fig. 1L). These results suggest that organoid-derived intestinal epithelial cells undergo turnover similar to that observed in the in vivo small intestine.

### Cell replication has a low impact on function

At Passage 12, organoid-derived intestinal epithelial cells exhibited a decrease in the proliferation marker Ki67-positive cells and a reduction in the cell proliferation rate (Fig. S2 and movie). To investigate the changes in gene expression with each passage, we performed comprehensive gene expression analysis (RNA-seq) on organoid-derived intestinal epithelial cells. Across Passages 6, 9, and 12, these cells showed minimal changes in overall mRNA expression (Fig. 2A). Additionally, in genes specific to the small intestine, there were few changes between passages, and organoid-derived intestinal epithelial cells from each passage clustered together, distinct from mesenchymal cells and cryopreserved human enterocytes (Fig. 2B). When comparing organoid-derived intestinal epithelial cells at different passages, the correlation coefficients (R^2^) were 0.855 (Passage 6 and 9) and 0.850 (Passage 6 and 12), indicating minimal changes between passages (Fig. 2C and D).

**Figure 2.**
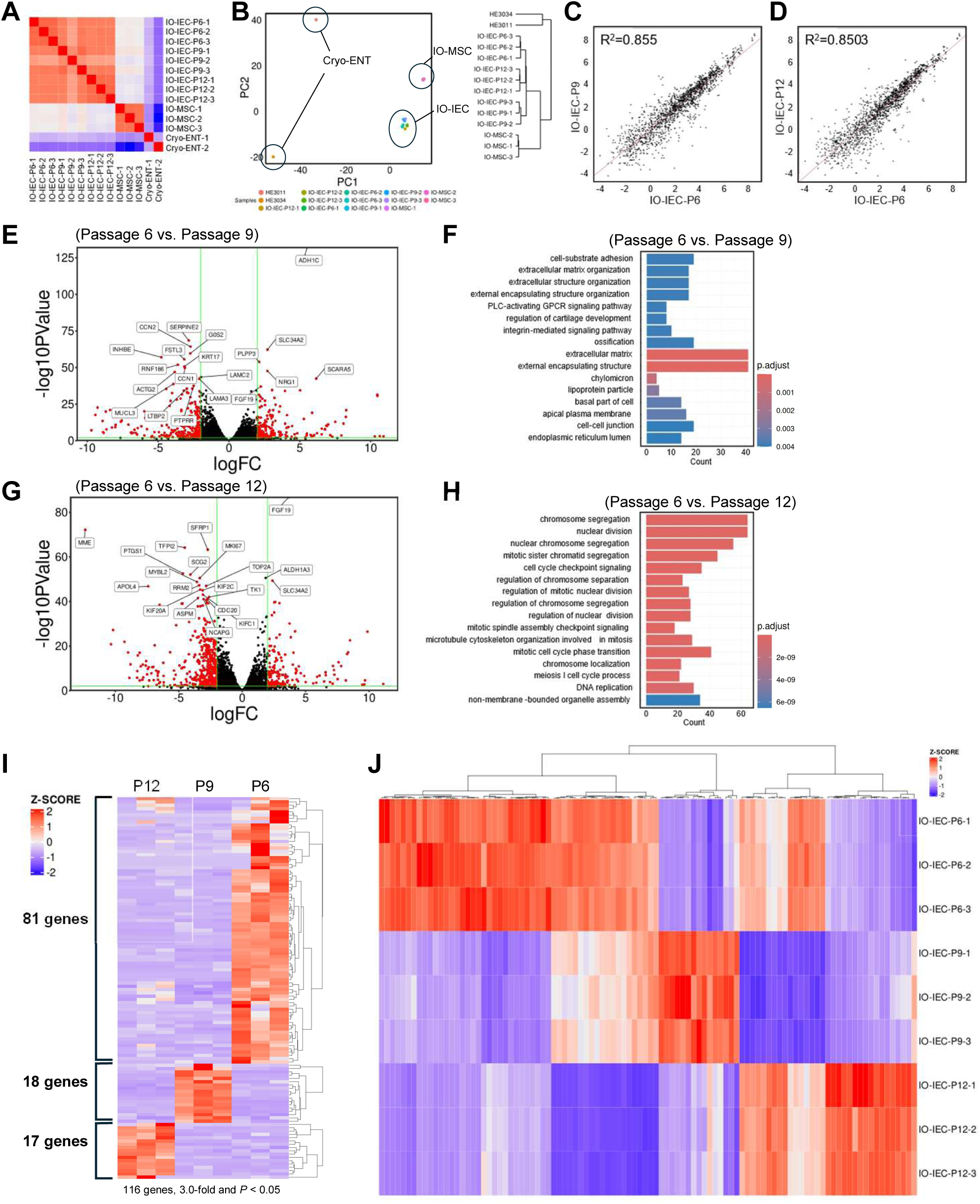
Changes in gene expression in organoid-derived intestinal epithelial cells. Intestinal organoid-derived intestinal epithelial cells are abbreviated as IO-IEC. Intestinal organoid-derived mesenchymal stromal cells are abbreviated as IO-MSC. Cryopreserved human enterocytes are abbreviated as Cryo-ENT. A. Correlation heatmap of organoid-derived intestinal epithelial cells, intestinal organoid-derived mesenchymal stromal cells, and cryopreserved human enterocytes. B. PCA map with hierarchical clustering of the small intestine features gene expression from organoid-derived intestinal epithelial cells, intestinal organoid-derived mesenchymal stromal cells, and cryopreserved human enterocytes. C. A scatter diagram of the small intestine features gene expression from organoid-derived intestinal epithelial cells Passage 6 and 9. D. A scatter diagram of the small intestine features gene expression from organoid-derived intestinal epithelial cells Passage 6 and 12. E. Volcano plot of intestinal organoid-derived intestinal epithelial cells Passage 6 and 9. F. GO enrichment analysis of intestinal organoid-derived intestinal epithelial cells Passage 6 and 9. G. Volcano plot of intestinal organoid-derived intestinal epithelial cells Passage 6 and 12. H. GO enrichment analysis of intestinal organoid-derived intestinal epithelial cells Passage 6 and 9. I. Heat map of differentially expressed in Intestinal organoid-derived intestinal epithelial cells Passage 6, 9, and 12. J. Heat map of characteristically expression genes in intestinal organoid-derived intestinal epithelial cells Passage 6, 9, and 12. This result was calculated using multiple comparisons.

To identify differentially expressed genes at Passage 9 and 12, compared to Passage 6, we conducted differential expression gene (DEG) analysis on organoid-derived intestinal epithelial cells. The study revealed that 3,231 genes were significantly altered, with 202 genes showing increased expression and 3,029 genes showing decreased expression at Passage 9 (Fig. 2E). The top 20 genes with the most significant changes were depicted in a volcano plot. Gene Ontology (GO) analysis showed that in organoid-derived intestinal epithelial cells at Passage 9, there were slight changes in gene expression related to extracellular matrix and external encapsulating structure pathways (Fig. 2F).

At Passage 12, organoid-derived intestinal epithelial cells exhibited significant alterations in 2,899 genes, with 168 genes showing increased expression and 2,731 genes showing decreased expression (Fig. 2G). The top 20 genes with the most significant changes were represented in a volcano plot. GO analysis indicated that organoid-derived intestinal epithelial cells at Passage 12 of many pathways related to cell proliferation are changed (Fig. 2H). Organoid-derived intestinal epithelial cells at Passages 6, 9, and 12 uniquely expressed 81, 18, and 17 genes, respectively (Fig. 2I and S3). While organoid-derived intestinal epithelial cells at Passage 9 had many genes related to the extracellular matrix, Passage 12 cells had more genes associated with tumor suppression, cell metabolism, and immune response.

Furthermore, multi-group comparison analysis across passages identified genes with a negative correlation between passage number and gene expression levels (Fig. 2J and S4). Many of these genes were involved in cell proliferation. These results indicate that the expression of genes related to cell proliferation in organoid-derived intestinal epithelial cells decreases with each passage, significantly affecting the cell proliferation rate at Passage 12 (Fig. 1K and 1L). The minimal changes in the expression of genes involved in intestinal epithelial cell functions (Fig. S5 and S6) suggest that cell functions are maintained despite repeated passaging.

### Barrier Function of Organoid–Derived Intestinal Epithelial Cells is Maintained Overtime

Intestinal epithelial cells function as a barrier separating the external environment from the internal body, making barrier function one of the key indicators for evaluating the functionality of in vitro intestinal epithelial cells. We, therefore, performed immunostaining related to barrier function and measured barrier functionality in passaged organoid-derived intestinal epithelial cells to investigate changes over 24 days. Organoid-derived intestinal epithelial cells at Passage 6, 9, and 12 expressed ECAD and CDH17, which are involved in adherens junctions, and ZO-1 and Claudin-7, which are involved in tight junctions. Furthermore, the formation of adherens junctions and tight junctions was confirmed using transmission electron microscopy (TEM) (Fig. 3A-F).

**Figure 3.**
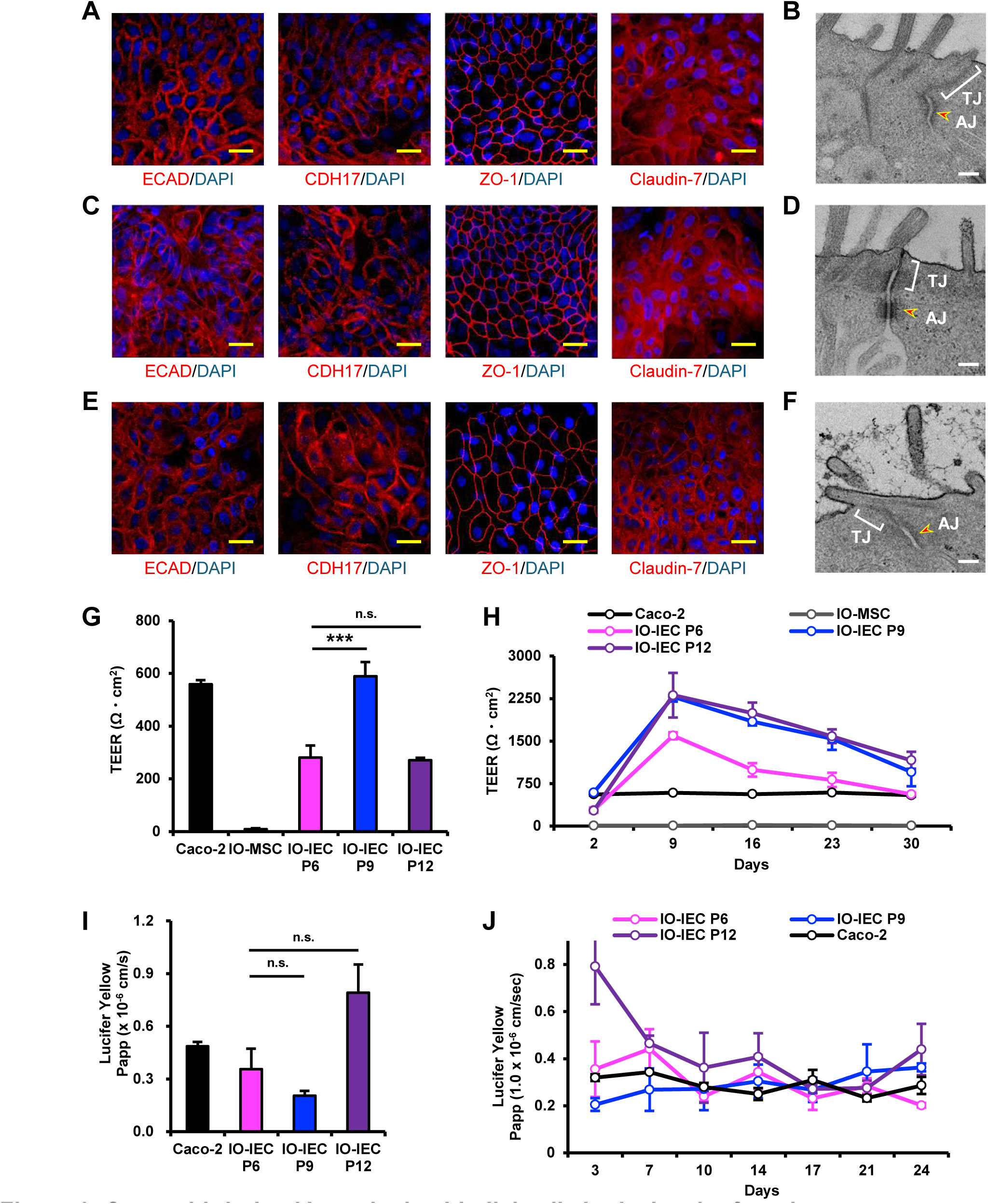
Organoid-derived intestinal epithelial cells show barrier function. Intestinal organoid-derived intestinal epithelial cells are abbreviated as IO-IEC. Intestinal organoid-derived mesenchymal stromal cells are abbreviated as IO-MSC. A. Immunocytochemistry of organoid-derived intestinal epithelial cells at Passage 6. Immunocytochemistry was performed with the antibody to barrier functions (ECAD, CDH17, ZO-1, and Claudin-7). Nuclei were stained with DAPI. Scale bars: 20 μm. B. Transmission electron microscopic analysis of epithelial lateral membrane of organoid-derived intestinal epithelial cells at Passage 6. Tight junction (TJ) and adherens junction (AJ, yellow arrow) were observed. Scale bar: 200 nm. C. Immunocytochemistry of organoid-derived intestinal epithelial cells at Passage 9. Immunocytochemistry was performed with the antibody to barrier functions (ECAD, CDH17, ZO-1, and Claudin-7). Nuclei were stained with DAPI. Scale bars: 20 μm. D. Transmission electron microscopic analysis of epithelial lateral membrane of organoid-derived intestinal epithelial cells at Passages 9. Tight junction (TJ) and adherens junction (AJ, yellow arrow) were observed. Scale bar: 200 nm. E. Immunocytochemistry of organoid-derived intestinal epithelial cells at Passage 12. Immunocytochemistry was performed with the antibody to barrier functions (ECAD, CDH17, ZO-1, and Claudin-7). Nuclei were stained with DAPI. Scale bars: 20 μm. F. Transmission electron microscopic analysis of epithelial lateral membrane of organoid-derived intestinal epithelial cells at Passage 12. Tight junction (TJ) and adherens junction (AJ, yellow arrow) were observed. Scale bar: 200 nm. G. Trans-epithelial electrical resistance measurements of organoid-derived intestinal epithelial cells at Passage 6, 9, and 12, intestinal organoid-derived mesenchymal stromal cells, and Caco-2 cells. Results are expressed as mean ± SD (n = 3 triplicate biological experiments). Statistical significance was determined using one-way ANOVA with Dunnett’s test compared to organoid-derived intestinal epithelial cells Passage 6. *** p < 0.001, ** p < 0.01, * p < 0.05. n.s.: not significant. H. Trans-epithelial electrical resistance measurements for 30 days cultured organoid-derived intestinal epithelial cells at Passage 6, 9, and 12, intestinal organoid-derived mesenchymal stromal cells, and Caco-2 cells. Results are expressed as mean ± SD (n = 3 triplicate biological experiments). I. Lucifer Yellow permeability test of organoid-derived intestinal epithelial cells at Passage 6, 9, and 12, and Caco-2 cells. Results are expressed as mean ± SD (n = 3 triplicate biological experiments). Statistical significance was determined using one-way ANOVA with Dunnett’s test compared to organoid-derived intestinal epithelial cells Passage 6. *** p < 0.001, ** p < 0.01, * p < 0.05. n.s.: not significant. J. Lucifer Yellow permeability test for 4 weeks cultured organoid-derived intestinal epithelial cells at Passage 6, 9, and 12, and Caco-2 cells. Results are expressed as mean ± SD (n = 3 triplicate biological experiments).

We conducted TEER measurements and Lucifer Yellow permeability assays to evaluate the barrier function. TEER measurements increased barrier function in organoid-derived intestinal epithelial cells at Passage 9 (Fig. 3G). The TEER values were 280 ± 46 Ω•cm^2^ for Passage 6, 589 ± 53 Ω•cm^2^ for Passage 9, and 271 ± 9 Ω•cm^2^ for Passage 12. The TEER values of organoid-derived intestinal epithelial cells fluctuated significantly over 30 days compared to Caco-2 cells (Fig. 3H and S7). The TEER values of organoid-derived intestinal epithelial cells reached their maximum on day 9 and subsequently decreased. At day 30, the TEER values were 563 ± 24 Ω•cm^2^ for Passage 6, 954 ± 251 Ω•cm^2^ for Passage 9, and 1161 ± 150 Ω•cm^2^ for Passage 12.

In the permeability assay using Lucifer Yellow, no significant differences were observed among organoid-derived intestinal epithelial cells at different Passage 6, 9, and 12 (Fig. 3I). The Papp values were 0.36 ± 0.1 × 10^−6^ cm/s for Passage 6, 0.20 ± 0.0 × 10^−6^ cm/s for Passage 9, and 0.79 ± 0.2 × 10^−6^ cm/s for Passage 12. Unlike the TEER assays, the Papp values remained constant over 24 days (Fig. 3J and S8). The minimal variation in Papp values indicates that the barrier function of organoid-derived intestinal epithelial cells is relatively stable. In contrast, the significant fluctuations in TEER values may be because organoid-derived intestinal epithelial cells encompass a variety of epithelial cell types compared to Caco-2 cells^15,18–22^.

### Transporter Function of Organoid–Derived Intestinal Epithelial Cells is Maintained Overtime

Intestinal epithelial cells permeate and efflux compounds, playing crucial roles in their absorption, distribution, metabolism, and excretion. Transporters are involved in transporting these compounds, and intestinal epithelial cells express a variety of transporters to handle the transport of diverse compounds^23–25^. Experiments using the small intestine are conducted to predict the pharmacokinetics and bioavailability of compounds and evaluate their toxicity and safety. This study assessed passaged organoid-derived intestinal epithelial cells by performing protein staining for P-gp, BCRP, and PEPT1 transporters. The mRNA expression levels of P-gp, BCRP, and PEPT1 in organoid-derived intestinal epithelial cells were equivalent to or exceeded those of cryopreserved human enterocytes but decreased with increasing passages (Fig. 4A). After four weeks of continuous culture, the expression of BCRP and PEPT1 tended to decline. In contrast, the expression of P-gp increased by 1.5- to 2.0-fold (Fig. 4B and S9). In passaged organoid-derived intestinal epithelial cells, the significant transporters P-gp, BCRP, and PEPT1 were expressed and localized to the apical membrane (Fig. 4C-E).

**Figure 4.**
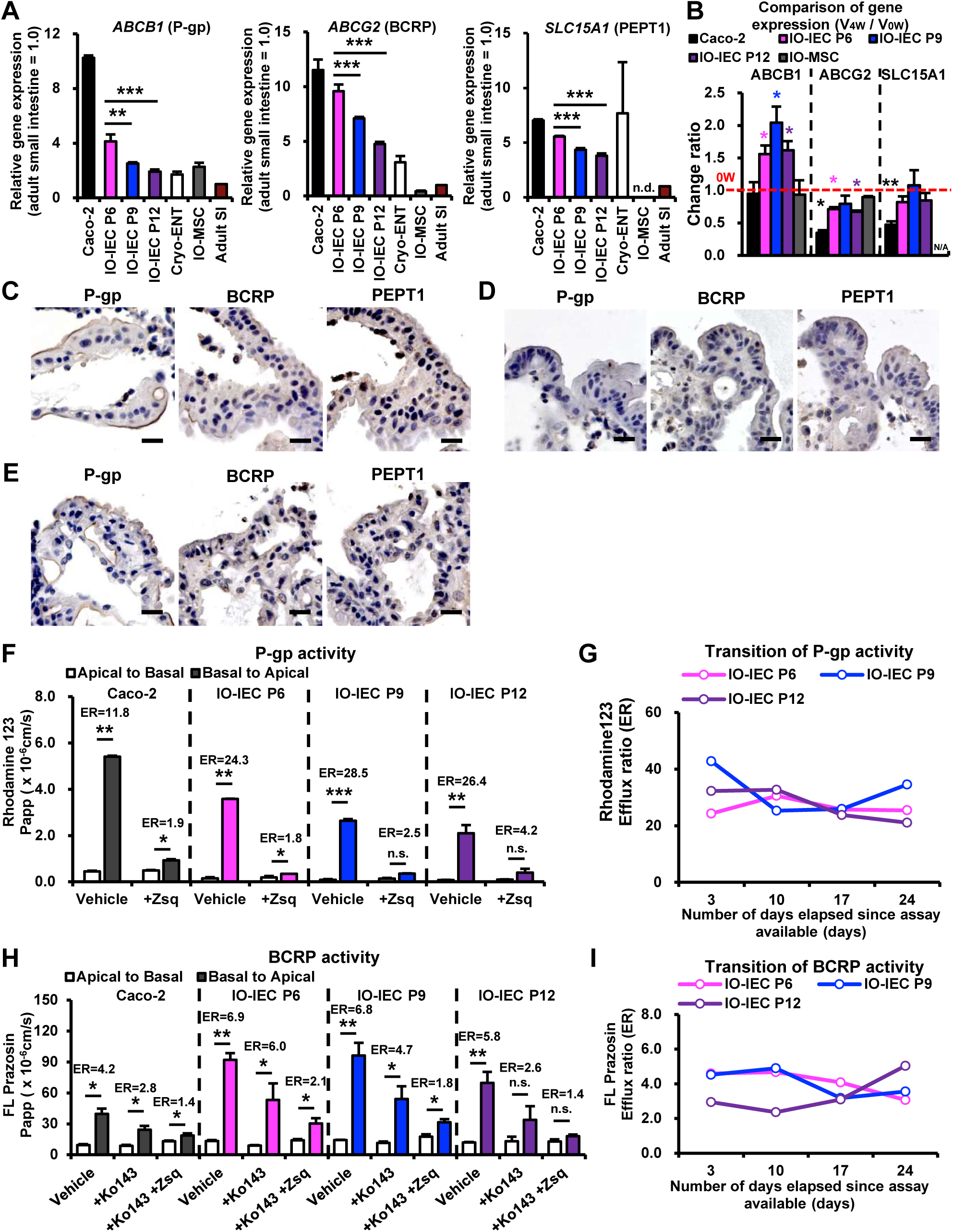
Organoid-derived intestinal epithelial cells exhibit transport capability. Intestinal organoid-derived intestinal epithelial cells are abbreviated as IO-IEC. Intestinal organoid-derived mesenchymal stromal cells are abbreviated as IO-MSC. Cryopreserved human enterocytes are abbreviated as Cryo-ENT. Adult small intestine is abbreviated as Adult SI. A. P-gp, BCRP, and PEPT1 gene expressions by qRT-PCR analysis in organoid-derived intestinal epithelial cells at Passage 6, 9, and 12, intestinal organoid-derived mesenchymal stromal cells, and Caco-2 cells in transwells. The expression level of adult small intestine whole tissue was set at 1.0. Results are expressed as mean ± SD (n = 3 triplicate biological experiments). Statistical significance was determined using one-way ANOVA with Dunnett’s test compared to organoid-derived intestinal epithelial cells Passage 6. *** p < 0.001, ** p < 0.01, * p < 0.05. n.s.: not significant. n.d.: not detected. B. Comparison of P-gp, BCRP, and PEPT1 gene expression in each cell at 0 and 4 weeks. The change ratio of gene expression was derived by dividing the relative gene expression values of 0 weeks (0W) and 4 weeks (4W). N/A: not available. C. Immunohistochemistry of intestinal epithelial cells at Passage 6 was performed with antibodies to P-gp, BCRP, and PEPT1. Scale bars: 20 μm. D. Immunohistochemistry of organoid-derived intestinal epithelial cells at Passage 9 was performed with antibodies to P-gp, BCRP, and PEPT1. Scale bars: 20 μm. E. Immunohistochemistry of organoid-derived intestinal epithelial cells at Passage 12 was performed with antibodies to P-gp, BCRP, and PEPT1. Scale bars: 20 μm. F. Permeability test with Rhodamine 123 in organoid-derived intestinal epithelial cells at Passage 6, 9, and 12, and Caco-2 cells. The efflux ratio (ER) for Rhodamine 123 was derived from the Papp values associated with basal-to-apical transport (B to A) and apical-to-basal transport (A to B). Results are expressed as mean ± SD (n = 3 triplicate biological experiments). Statistical significance was determined using Student’s t-test; *** p < 0.001, ** p < 0.01, * p < 0.05. Rhodamine 123: P-gp substrate. Zsq: zosuquidar, P-gp inhibitor. G. The efflux ratio (ER) for Rhodamine 123 for 4 weeks cultured organoid-derived intestinal epithelial cells at Passage 6, 9, and 12. H. Permeability test with FL Prazosin in organoid-derived intestinal epithelial cells at Passage 6, 9, and 12, and Caco-2 cells. The efflux ratio (ER) for FL Prazosin was derived from the Papp values associated with basal-to-apical transport (B to A) and apical-to-basal transport (A to B). Results are expressed as mean ± SD (n = 3 triplicate biological experiments). Statistical significance was determined using Student’s t-test; *** p < 0.001, ** p < 0.01, * p < 0.05. FL Prazosin: BCRP and P-gp substrate. Ko143: BCRP inhibitor. Zsq: zosuquidar, P-gp inhibitor. I. The efflux ratio (ER) for FL Prazosin for 4 weeks cultured organoid-derived intestinal epithelial cells at Passage 6, 9, and 12.

Next, to evaluate the transport activities of P-gp and BCRP, we conducted permeability assays using substrates and inhibitors. The transport activity was assessed by calculating the Efflux Ratio (ER) from the permeability coefficients, where 2.0 or higher indicates asymmetric transport. In the transport assay using Rhodamine 123, a substrate for P-gp, the ER value for Passage 6 was 24.3, which decreased to 1.8 upon adding the inhibitor Zosuquidar. For Passage 9, the ER value was 28.5, which decreased to 2.5 with the addition of Zosuquidar. At Passage 12, the ER value was 26.4, decreasing to 4.2 with the addition of the inhibitor (Fig. 4F). Furthermore, during four weeks of measuring P-gp activity, the ER values for Passage 6 ranged from 24.3 to 30.5, Passage 9 from 25.3 to 42.8, and Passage 12 from 21.1 to 32.7 (Fig. 4G and S10). Interestingly, the results of the transporter activity did not correlate with the mRNA expression levels, indicating that mRNA expression does not necessarily reflect transporter activity (Fig. 4A and 4G).

In the transport assay using FL-Prazosin, a substrate for both BCRP and P-gp, the ER values for Passage 6 were 6.9, which decreased to 6.0 with the addition of the BCRP inhibitor Ko 143 and further decreased to 2.1 with the addition of both Ko 143 and the P-gp inhibitor Zosuquidar. For Passage 9, the ER value was 6.8, which decreased to 4.7 with Ko 143 and 1.8 with both inhibitors. At Passage 12, the ER value was 5.8, decreasing to 2.6 with Ko 143 and further to 1.4 with both Ko 143 and Zosuquidar (Fig. 4H). During four weeks of measuring BCRP activity, the ER values for Passage 6 ranged from 3.1 to 4.7, Passage 9 from 3.2 to 4.9, and Passage 12 from 2.4 to 5.0 (Fig. 4I and S11). These results demonstrate that organoid-derived intestinal epithelial cells at Passage 6, 9, and 12 possess the activity of P-gp and BCRP and maintain this functionality over extended periods. Additionally, the transport activities were effectively inhibited by known inhibitors, making these cells suitable for predicting and evaluating intestinal behavior in drug discovery research.

### Organoid–derived Intestinal Epithelial Cells have Drug Metabolic Capacity

We extended our investigation to passaged organoid-derived intestinal epithelial cells, assessing the expression and activity of CYPs, including CYP3A4 and CES enzymes. We performed immunostaining for CYP3A5 and CES2, as well as measured CYPs (CYP1A1, 2B6, 2C9, 2C19, 2D6, 3A4) and CES expression levels and enzymatic activities in passaged organoid-derived intestinal epithelial cells. The mRNA expression levels of CYP1A1, 2C9, 2C19, and 3A4 in organoid-derived intestinal epithelial cells were equivalent to or exceeded those of cryopreserved human enterocytes (Fig. 5A). In contrast, the expression of CYP1A1 in organoid-derived intestinal epithelial cells was exceptionally high, approximately 900-fold higher than that in cryopreserved human enterocytes at Passage 6. CYP2B6 and CYP2D6 in organoid-derived intestinal epithelial cells exhibited lower expression than cryopreserved human enterocytes, with CYP2D6 being particularly low at approximately 1/80 the expression level.

**Figure 5.**
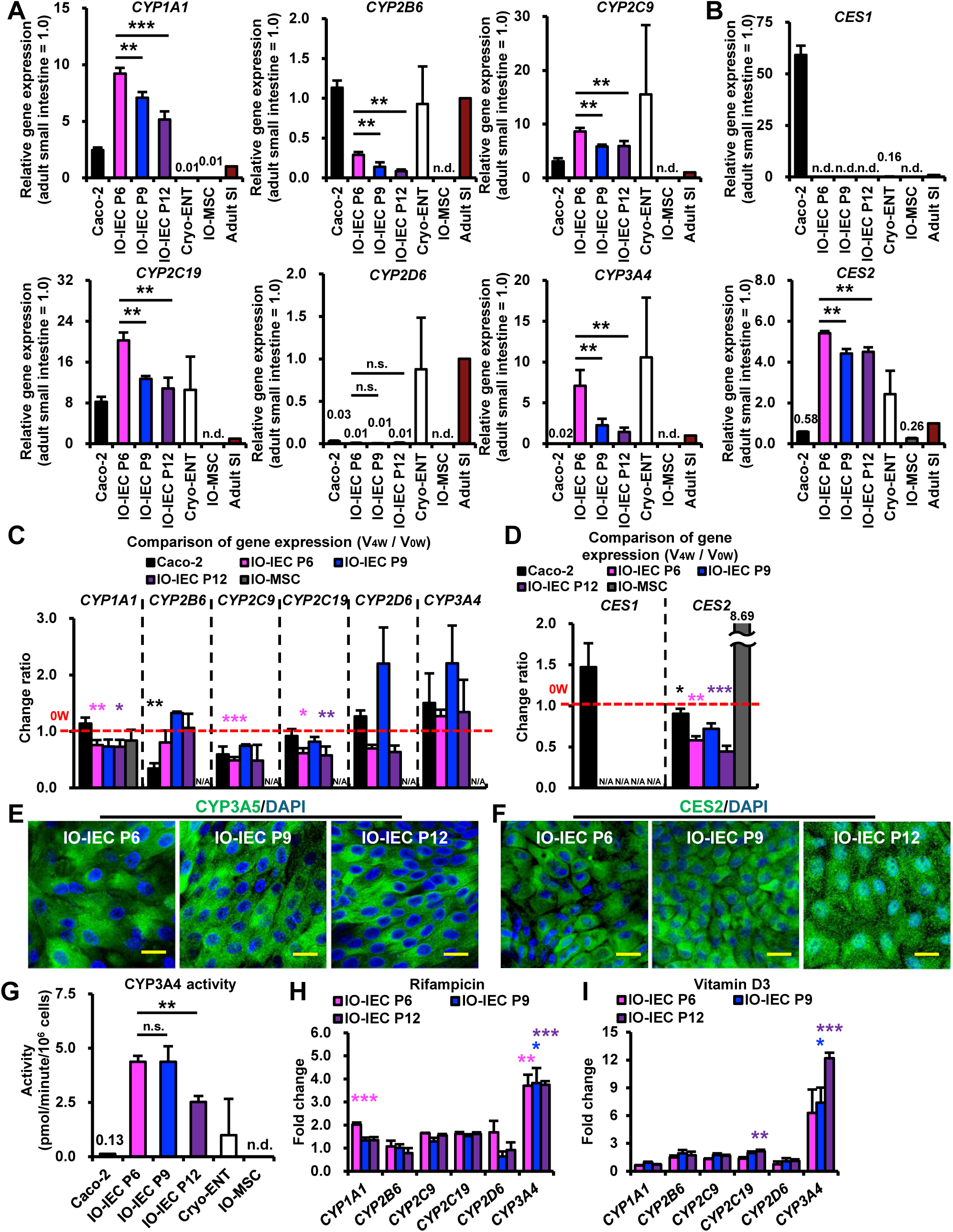
Organoid-derived intestinal epithelial cells have drug-metabolizing ability. Intestinal organoid-derived intestinal epithelial cells are abbreviated as IO-IEC. Intestinal organoid-derived mesenchymal stromal cells are abbreviated as IO-MSC. Cryopreserved human enterocytes are abbreviated as Cryo-ENT. Adult small intestine is abbreviated as Adult SI. A. Expression of cytochrome P450 genes (CYP1A1, CYP2B6, CYP2C9, CYP2C19, CYP2D6, and CYP3A4) in Caco-2 cells, organoid-derived intestinal epithelial cells, cryopreserved human enterocytes, and intestinal organoid-derived mesenchymal stromal cells. The expression levels of adult small intestine whole tissue were set to 1.0. Expression levels were calculated from the results of independent (biological) triplicate experiments. Results are expressed as mean ± SD (n = 3). Statistical significance was determined using one-way ANOVA with Dunnett’s test compared to organoid-derived intestinal epithelial cells Passage 6. *** p < 0.001, ** p < 0.01, * p < 0.05. n.s.: not significant. n.d.: not detected. B. Expression of carboxylesterase (CES1 and CES2) in Caco-2 cells, organoid-derived intestinal epithelial cells, cryopreserved human enterocytes, and intestinal organoid-derived mesenchymal stromal cells. The expression levels of adult small intestine whole tissue were set to 1.0. Expression levels were calculated from the results of independent (biological) triplicate experiments. Results are expressed as mean ± SD (n = 3). Statistical significance was determined using one-way ANOVA with Dunnett’s test compared to organoid-derived intestinal epithelial cells Passage 6. *** p < 0.001, ** p < 0.01, * p < 0.05. n.s.: not significant. n.d.: not detected. C. Comparison of cytochrome P450 (CYP1A1, CYP2B6, CYP2C9, CYP2C19, CYP2D6, and CYP3A4) gene expression in each cell at 0 and 4 weeks. The change ratio of gene expression was derived by dividing the relative gene expression values of 0 weeks (0W) and 4 weeks (4W). N/A: not available. D. Comparison of carboxylesterase (CES1 and CES2) gene expression in each cell at 0 and 4 weeks. The change ratio of gene expression was derived by dividing the relative gene expression values of 0 weeks (0W) and 4 weeks (4W). N/A: not available. E. Immunocytochemistry of organoid-derived intestinal epithelial cells at Passage 6, 9, and 12 with the antibody to CYP3A5. Nuclei were stained with DAPI. Scale bars: 20 μm. F. Immunocytochemistry of organoid-derived intestinal epithelial cells at Passage 6, 9, and 12 with the antibody to CES2. Nuclei were stained with DAPI. Scale bars: 20 μm. G. Measure CYP3A4 activity in Caco-2 cells, organoid-derived intestinal epithelial cells, and intestinal organoid-derived mesenchymal stromal cells. Results are expressed as mean ± SD (n = 3 triplicate biological experiments, cryopreserved human enterocytes n = 6). Statistical significance was determined using one-way ANOVA with Dunnett’s test compared to organoid-derived intestinal epithelial cells Passage 6. *** p < 0.001, ** p < 0.01, * p < 0.05. n.s.: not significant. n.d.: not detected. H. Induction of the cytochrome P450 genes with exposure to rifampicin (Rif). Results are expressed as mean ± SD (n = 3). The expression level of each gene without any treatment (DMSO) was set to 1.0. Expression levels were calculated from the results of independent (biological) triplicate experiments. Statistical significance was determined using Student’s t-test: *** Fold > 2, p < 0.001, ** Fold > 2, p < 0.01, * Fold > 2, p < 0.05. I. Induction of the cytochrome P450 genes with vitamin D (VD3) exposure. Results are expressed as mean ± SD (n = 3). The expression level of each gene without any treatment (DMSO) was set to 1.0. Expression levels were calculated from the results of independent (biological) triplicate experiments. Statistical significance was determined using Student’s t-test: *** Fold > 2, p < 0.001, ** Fold > 2, p < 0.01, * Fold > 2, p < 0.05.

Carboxylesterase (CES2) gene expression in organoid-derived intestinal epithelial cells was comparable to that in cryopreserved human enterocytes. There was no expression of the liver-type CES1 (Fig. 5B). Although the expression of CYPs in organoid-derived intestinal epithelial cells tended to decrease with increasing passages, the expression of CES2 was maintained across passages (Fig. 5A and 5B). Furthermore, after four weeks of culture, CYP1A1, 2C9, 2C19, and CES2 expression levels in organoid-derived intestinal epithelial cells decreased. Still, except for CYP2C9, the levels remained equivalent to or exceeded those of cryopreserved human enterocytes (Fig. 5C, 5D, S12, and S13). Immunostaining revealed that organoid-derived intestinal epithelial cells at all passages were positive for CYP3A5 and CES2 (Fig. 5E and 5F).

In terms of CYP metabolic activity, organoid-derived intestinal epithelial cells exhibited CYP2B6, 2C9, 2C19, and 2D6 activities comparable to Caco-2 cells, while CYP3A4 activity was approximately 34-fold higher (Fig. 5G and S12). This CYP3A4 activity was high, about 4-fold that of cryopreserved human enterocytes. Although CYP3A4 activity in organoid-derived intestinal epithelial cells decreased at Passage 12, the activities of other CYPs remained unchanged (Fig. 5G and S14). Moreover, in all passages of organoid-derived intestinal epithelial cells, the expression of CYPs was induced by vitamin D3, omeprazole, rifampicin, dexamethasone, and phenobarbital (Fig. 5H, 5I, and S15). Notably, Phenobarbital-induced CYP2B6 expression was not observed in organoid-derived intestinal epithelial cells. This is likely due to the absence of Constitutive Androstane Receptor (CAR) expression in these cells, similar to the small intestine^26^.

Finally, compounds such as Naproxen and Acetaminophen are toxic to small intestinal epithelial cells^27,28^. When naproxen and acetaminophen were administered to Caco-2 and organoid-derived intestinal epithelial cells, both cell types exhibited cytotoxicity (Fig. 6). Organoid-derived intestinal epithelial cells showed cytotoxicity starting at 1.25 μM. It exhibited dose-dependent cytotoxicity (Fig. 6A, 6D, and 6E). In contrast, Caco-2 cells only showed significant cytotoxicity at a high concentration of 10 μM (Fig. 6A). Acetaminophen exhibited weaker toxicity in organoid-derived intestinal epithelial cells due to the low expression of CYP2E1, which metabolizes it. Minimal toxicity started at 25 μM, with significant cell death observed on the second day after administration (Fig. 6A, 6F, and 6G).

**Figure 6.**
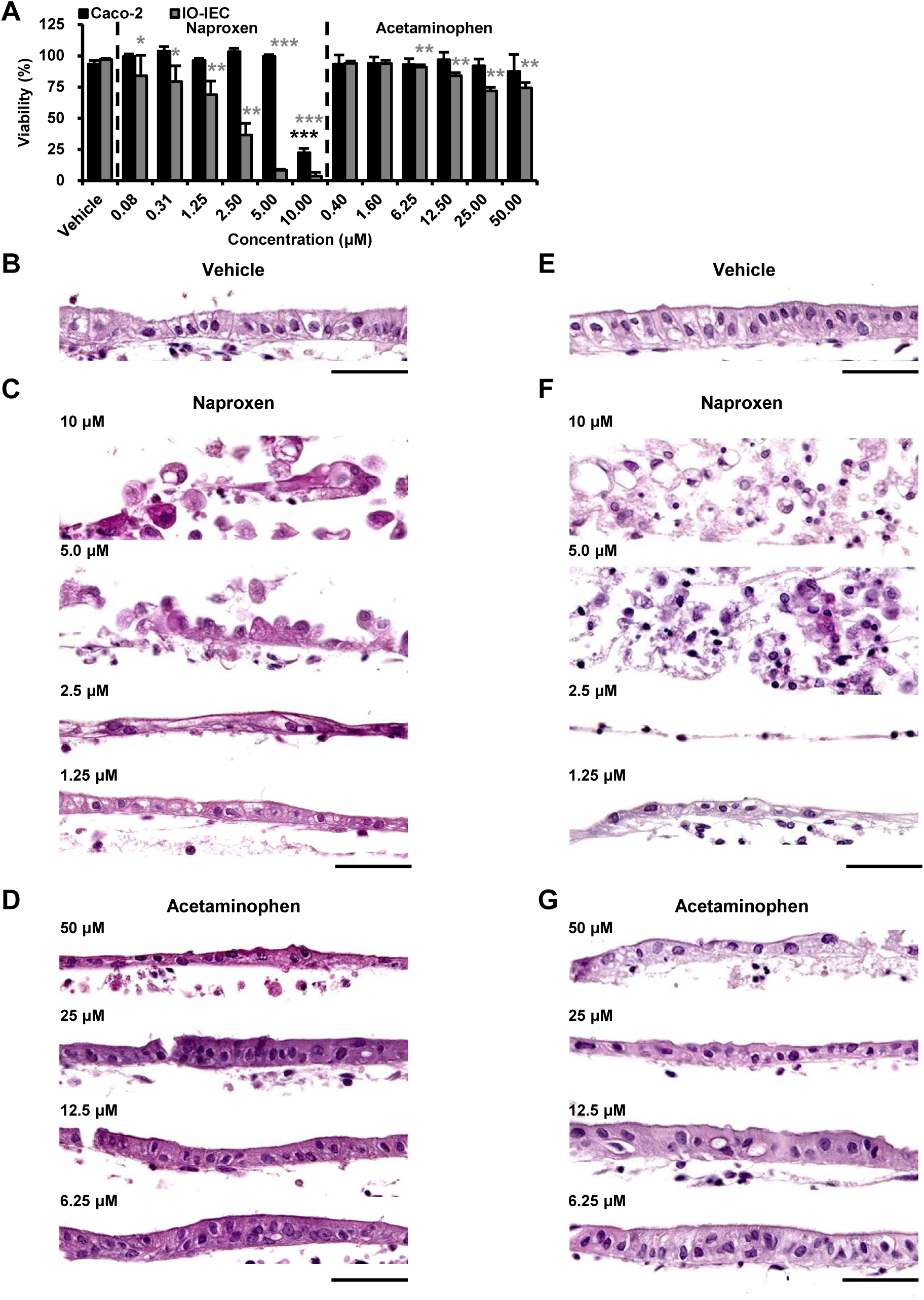
Naproxen and acetaminophen toxicity in organoid-derived intestinal epithelial cells. Intestinal organoid-derived intestinal epithelial cells are abbreviated as IO-IEC. A. Viability of organoid-derived intestinal epithelial cells and Caco-2 cells exposed to naproxen and acetaminophen for 1 day. We determined cell viability using the WST-1 Cell Proliferation Assay System. B. Histology of organoid-derived intestinal epithelial cells exposed to DMSO for 1 day. Scale bar: 50 μm. C. Histology of organoid-derived intestinal epithelial cells exposed to respective naproxen concentrations for 1 day. Scale bar: 50 μm. D. Histology of organoid-derived intestinal epithelial cells exposed to respective concentrations of acetaminophen for 1 day. Scale bar: 50 μm. E. Histology of organoid-derived intestinal epithelial cells exposed to DMSO for 2 days. Scale bar: 50 μm. F. Histology of organoid-derived intestinal epithelial cells exposed to respective naproxen concentrations for 2 days. Scale bar: 50 μm. G. Histology of organoid-derived intestinal epithelial cells exposed to respective concentrations of acetaminophen for 2 days. Scale bar: 50 μm.

## 3. Discussion

As shown in this study, passage-independent cell characteristics of organoid-derived intestinal epithelial cells imply that organoid-derived intestinal epithelial cells up to Passage 9 are appropriate for studies such as drug discovery research. Organoid-derived intestinal epithelial cells at Passage 9 have undergone 18 divisions and cover an area exceeding 400 m^2^, indicating that highly functional small intestinal epithelial cells can be stably supplied. The function of organoid-derived intestinal epithelial cells is to maintain them over time, allowing their application in administration tests. This extends the usable period in multi-organ devices such as microphysiological systems and organ-on-chip technologies^29–31^. Likewise, organoid-derived intestinal epithelial cells’ drug metabolism and transport capabilities, comparable to those of Caco-2 cells and cryopreserved human enterocytes, indicate their utility in drug discovery research.

CYP3A4 and CYP3A5 are highly expressed in both the liver and the intestine, and intestinal CYP3A4 and CYP3A5 are responsible for the initial metabolism of orally administered drugs^32^. This makes them extremely important for predicting bioavailability and pharmacokinetics. CYP3A4 and CYP3A5 metabolize both midazolam and testosterone^33–39^. The differences in the midazolam and testosterone metabolism assays in our study between organoid-derived intestinal epithelial cells and human enterocytes are likely attributable to the distinct ratios of CYP3A4, CYP3A5, and CYP3A7. The expression of CYP2D6 in organoid-derived intestinal epithelial cells was significantly lower than in cryopreserved human enterocytes and adult small intestine, likely due to the absence of bile acids in this system. This finding is consistent with bile acids enhancing CYP2D6 expression in the intestine^40,41^. Therefore, co-culturing organoid-derived intestinal epithelial cells with hepatocytes or adding bile acids to this system may induce CYP2D6 expression.

According to guidance from the U.S. Food and Drug Administration (FDA), pharmaceutical candidates are required to be evaluated as substrates of P-glycoprotein (P-gp) and Breast Cancer Resistance Protein (BCRP) at early stages of drug development (https://www.fda.gov/drugs/guidance-compliance-regulatory-information/guidances-drugs). Organoid-derived intestinal epithelial cells exhibited activity for both P-gp and BCRP, and their inhibition by known inhibitors such as zosuquidar and Ko143 suggests that these cells fulfill the FDA’s evaluation criteria. Furthermore, organoid-derived intestinal epithelial cells displayed a similar expression pattern of carboxylesterases (CES) to cryopreserved human enterocytes, as well as potent cytotoxicity with the CYP2C9 substrate naproxen and weak cytotoxicity with the CYP2E1 substrate acetaminophen^27,28^. These findings, reflecting drug metabolism behaviors akin to those in the human intestine, suggest that organoid-derived intestinal epithelial cells more accurately represent the functional properties of intestinal epithelial cells than Caco-2 cells. Therefore, they present an attractive model for evaluating the pharmacokinetics of drugs in the human small intestine.

## 4. Conclusion

We demonstrated that organoid-derived intestinal epithelial cells exhibit high stability, maintaining their functionality as small intestinal epithelial cells through the optimization of cell division. In drug discovery applications, their barrier function, transporter activity, and drug metabolism capacity are comparable to those of the widely used Caco-2 cells and cryopreserved human enterocytes. Consequently, organoid-derived intestinal epithelial cells represent a valuable novel model for drug development.

## Declarations

## Ethics approval and consent to participate

The Institutional Review Board approved all experiments involving human cells and tissue handling at the National Center for Child Health and Development (2021-178). In compliance with the Declaration of Helsinki, informed consent was obtained from all tissues and cell donors. When the donors were under 18, informed consent was obtained from parents. Human cells in this study were utilized in full compliance with the Ethical Guidelines for Medical and Health Research Involving Human Subjects (Ministry of Health, Labor, and Welfare (MHLW), Japan; Ministry of Education, Culture, Sports, Science and Technology (MEXT, Japan) and performed in full compliance with the Ethical Guidelines for Clinical Studies (Ministry of Health, Labor, and Welfare, Japan). The cells were banked after approval of the Institutional Review Board at the National Institute of Biomedical Innovation (May 9, 2006). The derivation and cultivation of human embryonic stem cell (hESC) lines were performed in full compliance with “the Guidelines for Derivation and Distribution of Human Embryonic Stem Cells (Notification of MEXT, No. 156 of August 21, 2009; Notification of MEXT, No. 86 of May 20, 2010) and “the Guidelines for Utilization of Human Embryonic Stem Cells (Notification of MEXT, No. 157 of August 21, 2009; Notification of MEXT, No. 87 of May 20, 2010)”. All procedures of animal experiments were approved by the Institutional Animal Care and Use Committee in the National Center for Child Health and Development, based on the basic guidelines for the conduct of animal experiments in implementing agencies under the jurisdiction of the Ministry of Health, Labour and Welfare (Notification of MHLW, No. 0220-1 of February 20, 2015). The Institutional Animal Care and Use Committee of the National Center for Child Health and Development approved the protocols of the animal experiments (approval number: A2003-002-C20-M01, title: Research on regenerative medicine and cell medicine using mesenchymal stem cells, iPS cells, and ESCs, and on toxicity testing). Animal experiments were performed according to protocols approved by the Institutional Animal Care and Use Committee of the National Research Institute for Child Health and Development.

## Consent for publication

Not applicable.

## Availability of data and material

The datasets and cells used during the current study are available from the corresponding author upon reasonable request.

## Authors’ contributions

CJ, SK, SH, SI, and AU designed the experiments. CJ, SK, SH, HK, and LT performed the experiments. CJ, SK, SH, and TKa analyzed data. HA, SI, and AU contributed to the reagents, tissues, and analysis tools. CJ, SK, SH, TKa, TKi, KN, SI, and AU discussed the data and manuscript. CJ, SK, and AU wrote this manuscript. All authors read and approved the final manuscript.

## Acknowledgments

We would like to express our sincere thanks to K. Miyado for fruitful discussion, to M. Ichinose for providing expert technical assistance, to C. Ketcham for English editing and proofreading, and to E. Suzuki and K. Saito for secretarial work.

## References

1. Zanger, U. M. & Schwab, M. Cytochrome P450 enzymes in drug metabolism: regulation of gene expression, enzyme activities, and impact of genetic variation. Pharmacol. Ther. 138, 103–141 (2013).

2. International Transporter Consortium et al. Membrane transporters in drug development. Nat. Rev. Drug Discov. 9, 215–236 (2010).

3. Arana, M. R., Tocchetti, G. N., Rigalli, J. P., Mottino, A. D. & Villanueva, S. S. M. Physiological and pathophysiological factors affecting the expression and activity of the drug transporter MRP2 in intestine. Impact on its function as membrane barrier. Pharmacol. Res. 109, 32–44 (2016).

4. Xue, Y., Ma, C., Hanna, I. & Pan, G. Intestinal transporter-associated drug absorption and toxicity. Adv. Exp. Med. Biol. 1141, 361–405 (2019).

5. Shi, Y. et al. Construction of the small intestine on molecular dynamics simulation and preliminary exploration of drug intestinal absorption prediction. Comput. Biol. Chem. 99, 107724 (2022).

6. Harwood, M. D. et al. In Vitro-In Vivo Extrapolation Scaling Factors for Intestinal P-Glycoprotein and Breast Cancer Resistance Protein: Part I: A Cross-Laboratory Comparison of Transporter-Protein Abundances and Relative Expression Factors in Human Intestine and Caco-2 Cells. Drug Metab. Dispos. 44, 297–307 (2016).

7. Sun, H., Chow, E. C., Liu, S., Du, Y. & Pang, K. S. The Caco-2 cell monolayer: usefulness and limitations. Expert Opin. Drug Metab. Toxicol. 4, 395–411 (2008).

8. Sato, T. et al. Single Lgr5 stem cells build crypt-villus structures in vitro without a mesenchymal niche. Nature 459, 262–265 (2009).

9. Spence, J. R. et al. Directed differentiation of human pluripotent stem cells into intestinal tissue in vitro. Nature 470, 105–109 (2011).

10. Wells, J. M. & Spence, J. R. How to make an intestine. Development 141, 752–760 (2014).

11. Ogaki, S., Shiraki, N., Kume, K. & Kume, S. Wnt and Notch signals guide embryonic stem cell differentiation into the intestinal lineages. Stem Cells 31, 1086–1096 (2013).

12. Ozawa, T. et al. Generation of enterocyte-like cells from human induced pluripotent stem cells for drug absorption and metabolism studies in human small intestine. Sci. Rep. 5, 16479 (2015).

13. Iwao, T. et al. Differentiation of human induced pluripotent stem cells into functional enterocyte-like cells using a simple method. Drug Metab. Pharmacokinet. 29, 44–51 (2014).

14. Inui, T. et al. Functional intestinal monolayers from organoids derived from human iPS cells for drug discovery research. Stem Cell Res. Ther. 15, 57 (2024).

15. Chen, J. et al. Human intestinal organoid-derived PDGFRα + mesenchymal stroma enables proliferation and maintenance of LGR4 + epithelial stem cells. Stem Cell Res. Ther. 15, 16 (2024).

16. Nishino, K. et al. Epigenetic-scale comparison of human iPSCs generated by retrovirus, Sendai virus or episomal vectors. Regen Ther 9, 71–78 (2018).

17. Nishino, K. et al. DNA methylation dynamics in human induced pluripotent stem cells over time. PLoS Genet. 7, e1002085 (2011).

18. Uchida, H., et al. A xenogeneic-free system generating functional human gut organoids from pluripotent stem cells. JCI Insight 2, e86492 (2017).

19. Tsuruta, S. et al. Development of Human Gut Organoids With Resident Tissue Macrophages as a Model of Intestinal Immune Responses. Cell Mol Gastroenterol Hepatol 14, 726–729.e5 (2022).

20. Artursson, P. Cell cultures as models for drug absorption across the intestinal mucosa. Crit. Rev. Ther. Drug Carrier Syst. 8, 305–330 (1991).

21. Hilgendorf, C. et al. Caco-2 versus Caco-2/HT29-MTX co-cultured cell lines: permeabilities via diffusion, inside- and outside-directed carrier-mediated transport. J. Pharm. Sci. 89, 63–75 (2000).

22. Hidalgo, I. J. Cultured intestinal epithelial cell models. Pharm. Biotechnol. 8, 35–50 (1996).

23. Glaeser, H. et al. Intestinal drug transporter expression and the impact of grapefruit juice in humans. Clin. Pharmacol. Ther. 81, 362–370 (2007).

24. Fromm, M. F. Importance of P-glycoprotein for drug disposition in humans. Eur. J. Clin. Invest. 33 Suppl 2, 6–9 (2003).

25. Brandsch, M., Knütter, I. & Bosse-Doenecke, E. Pharmaceutical and pharmacological importance of peptide transporters. J. Pharm. Pharmacol. 60, 543–585 (2008).

26. Janssen, A. W. F. et al. Cytochrome P450 expression, induction and activity in human induced pluripotent stem cell-derived intestinal organoids and comparison with primary human intestinal epithelial cells and Caco-2 cells. Arch. Toxicol. 95, 907–922 (2021).

27. Tracy, T. S., Marra, C., Wrighton, S. A., Gonzalez, F. J. & Korzekwa, K. R. Involvement of multiple cytochrome P450 isoforms in naproxen O-demethylation. Eur. J. Clin. Pharmacol. 52, 293–298 (1997).

28. Laine, J. E., Auriola, S., Pasanen, M. & Juvonen, R. O. Acetaminophen bioactivation by human cytochrome P450 enzymes and animal microsomes. Xenobiotica 39, 11–21 (2009).

29. Kurniawan, D. A. et al. Gut-liver microphysiological systems revealed potential crosstalk mechanism modulating drug metabolism. PNAS Nexus 3, gae070 (2024).

30. Carvalho, M. R. et al. Gastrointestinal organs and organoids-on-a-chip: advances and translation into the clinics. Biofabrication 15, (2023).

31. Fedi, A. et al. In vitro models replicating the human intestinal epithelium for absorption and metabolism studies: A systematic review. J. Control. Release 335, 247–268 (2021).

32. Paine, M. F. et al. THE HUMAN INTESTINAL CYTOCHROME P450 ‘PIE’. Drug Metab. Dispos. 34, 880–886 (2006).

33. Hustert, E. et al. The genetic determinants of the CYP3A5 polymorphism. Pharmacogenetics 11, 773–779 (2001).

34. Kuehl, P. et al. Sequence diversity in CYP3A promoters and characterization of the genetic basis of polymorphic CYP3A5 expression. Nat. Genet. 27, 383–391 (2001).

35. Schuetz, J., Beach, D. L. & Guzelian, P. Selective expression of cytochrome P450 CYP3A mRNAs in embryonic and adult human liver. Pharmacogenetics 4, 11–20 (1994).

36. Greuet, J., Pichard, L., Bonfils, C., Domergue, J. & Maurel, P. The fetal specific gene CYP3A7 is inducible by rifampicin in adult human hepatocytes in primary culture. Biochem. Biophys. Res. Commun. 225, 689–694 (1996).

37. Huang, H. et al. Transcription factors and ncRNAs associated with CYP3A expression in human liver and small intestine assessed with weighted gene co-expression network analysis. Biomedicines 10, 3061 (2022).

38. de Wildt, S. N., Kearns, G. L., Leeder, J. S. & van den Anker, J. N. Cytochrome P450 3A: ontogeny and drug disposition: Ontogeny and Drug Disposition. Clin. Pharmacokinet. 37, 485–505 (1999).

39. Guo, Y. et al. Quantitative prediction of CYP3A4- and CYP3A5-mediated drug interactions. Clin. Pharmacol. Ther. 107, 246–256 (2020).

40. Yerra, V. G. & Drosatos, K. Specificity proteins (SP) and Krüppel-like factors (KLF) in liver physiology and pathology. Int. J. Mol. Sci. 24, (2023).

41. Pan, X., Ning, M. & Jeong, H. Transcriptional regulation of CYP2D6 expression. Drug Metab. Dispos. 45, 42–48 (2017).

42. FDA Guidances. U. S. Food and Drug Administration. https://www.fda.gov/drugs/guidance-compliance-regulatory-information/guidances-drugs. Accessed on January 9, 2025.

