## Supplementary Figure 1 for "Human pluripotent stem cell-derived intestinal epithelial cells maintain small intestine-specific functions over time, even with repeated cell division"

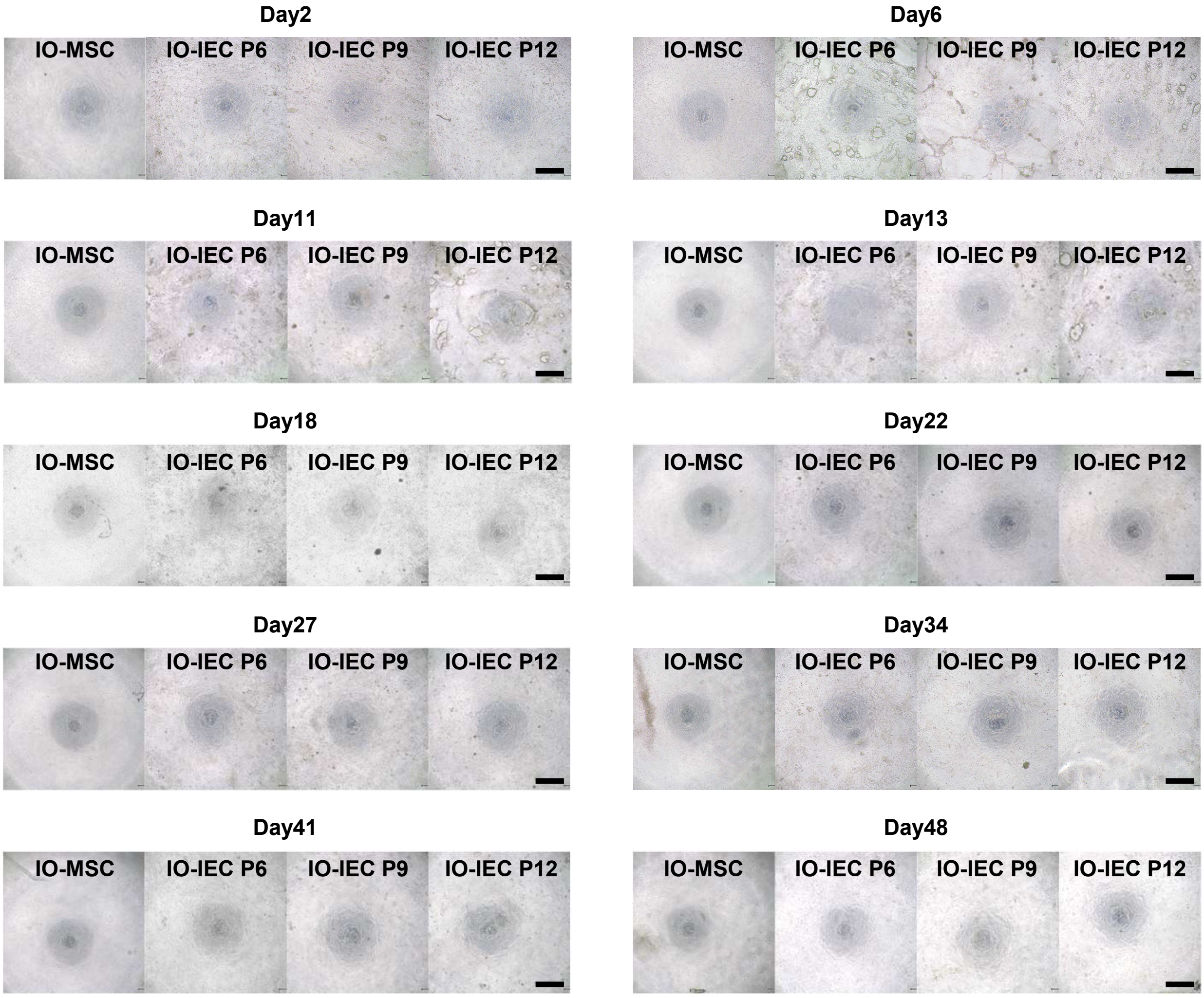


**Supplementary Figure 1. Phase-contrast micrograph of organoid-derived intestinal epithelial cells and intestinal organoid-derived mesenchymal stromal cells during proliferation**

**Supplementary Figure 1. Phase-contrast micrograph of organoid-derived intestinal epithelial cells and intestinal organoid-derived mesenchymal stromal cells during proliferation**

Intestinal organoid-derived intestinal epithelial cells are abbreviated as IO-IEC. Intestinal organoid-derived mesenchymal stromal cells are abbreviated as IO-MSC. Phase-contrast micrograph of organoid-derived intestinal epithelial cells (Passage 6, 9, and 12) and intestinal organoid-derived mesenchymal stromal cells for each timepoint. Scale bars: 500 μm.
