## Supplementary Figure 2 for "Human pluripotent stem cell-derived intestinal epithelial cells maintain small intestine-specific functions over time, even with repeated cell division"

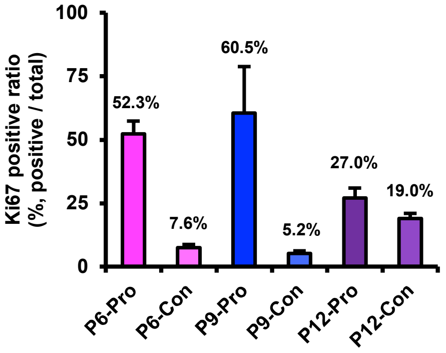


**Supplementary Figure 2. The change in Ki67 positivity in organoid-derived intestinal epithelial cells**

**Supplementary Figure 2. The change in Ki67 positivity in organoid-derived intestinal epithelial cells**

The state of proliferation is abbreviated as pro. The state of confluent is abbreviated as -con. The number of organoid-derived intestinal epithelial cells counted at Passage 6, 9, and 12: 4035, 3301 and 3779, respectively. Results are expressed as mean ± SD.
