## Supplementary Figure 3 for "Human pluripotent stem cell-derived intestinal epithelial cells maintain small intestine-specific functions over time, even with repeated cell division"

| Cell types | Gene symbol | | | | | | |
|---|---|---|---|---|---|---|---|
| **Passage 6-high**<br>**81 genes** | GALNT15 | IL11 | LINC00702 | MPP4 | DOCK2 | PTGS1 | RNF186 |
| | HOXA9 | TFPI2 | SERPINE1 | APOA4 | GREM1 | TRIM40 | SLFN11 |
| | LINC00607 | RELN | MMP3 | SLC28A1 | APOB | HOXA10 | IL21R |
| | SLA | PI16 | EGLN3-AS1 | SHROOM3 | ODAD2 | MME | C8orf34 |
| | TRBV30 | ZNF322P1 | FAM25A | C17orf78 | APLN | RIPOR3-AS1 | FGD2 |
| | NPY4R2 | ADAMTS8 | NEB | FMO3 | LINC01615 | CGB8 | FIBIN |
| | INSYN2B | SLCO5A1 | VNN1 | ARHGAP22 | HTR3A | ITGBL1 | PCDHGA12 |
| | SORL1-AS1 | IL16 | DIRAS2 | SLC8A1 | LINC02454 | HOXA13 | RFX8 |
| | LINC01182 | HOTTIP | SRI-AS1 | APBB1IP | DIAPH1 | NELL2 | IL37 |
| | GALNT6 | ANKRD31 | MEDAG | RASSF2 | SEPTIN4-AS1 | HOXA11-AS | TXK |
| | CTNS-AS1 | LINC01807 | LRRC56 | EGOT | LINC00648 | LINC02754 | NOX4 |
| | ITGAL | CCN4 | RPLP0P2 | TRPV2 | | | |
| **Passage 9-high**<br>**18 genes** | PALD1 | ADH4 | LAMP5 | SSTR2 | ADH6 | C1QL1 | OTC |
| | SOSTDC1 | ADH1C | EVA1A-AS | SCARA5 | GRIK1 | DUSP9 | HAS3 |
| | LGR5 | SYNPR | ACTL8 | IQSEC1 | | | |
| **Passage 12-high**<br>**17 genes** | PDZK1IP1 | KRT4 | SLURP2 | WBP1LP2 | SAA2-SAA4 | TRABD2B | CMKLR2 |
| | RASA4B | OLR1 | DSCAML1 | PRKN | LINC02925 | FGA | SPX |
| | NAV3 | DMBT1 | SHOC1 | | | | |

**Supplementary Figure 3. Highly expressed genes in organoid-derived intestinal epithelial cells at each passage**

**Supplementary Figure 3. Highly expressed genes in organoid-derived intestinal epithelial cells at each passage**

List of genes differentially expressed in Intestinal organoid-derived intestinal epithelial cells Passage 6, 9, and 12. These data correspond to the heat map in Figure. 2I.
