## Supplementary Figure 4 for "Human pluripotent stem cell-derived intestinal epithelial cells maintain small intestine-specific functions over time, even with repeated cell division"

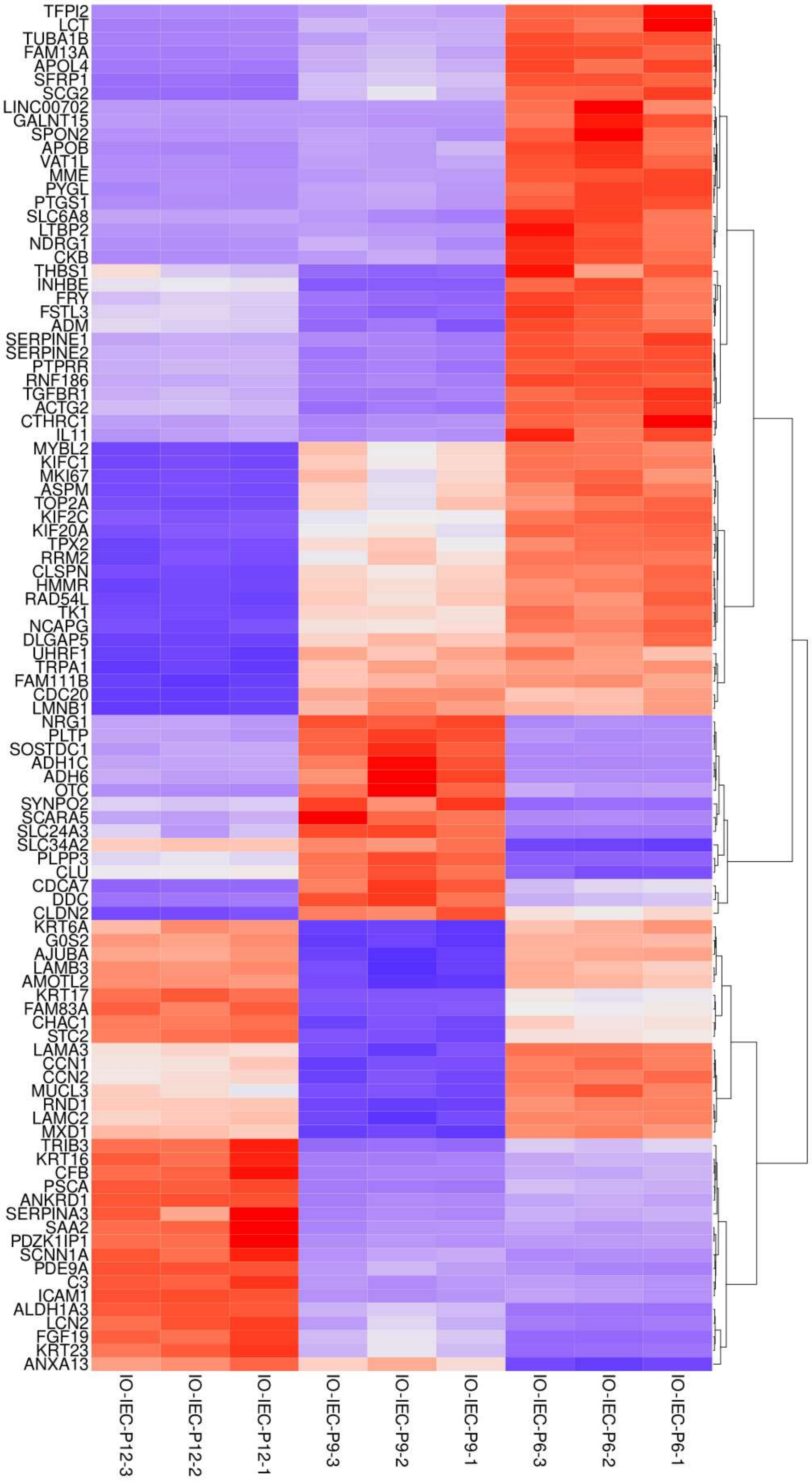


**Supplementary Figure 4. The featured genes in organoid-derived intestinal epithelial cells at each passage**

**Supplementary Figure 4. The featured genes in organoid-derived intestinal epithelial cells at each passage**

Intestinal organoid-derived intestinal epithelial cells are abbreviated as IO-IEC. List of featured expression genes in intestinal organoid-derived intestinal epithelial cells Passage 6, 9, and 12. This result was calculated using multiple comparisons. This data corresponds to the heat map in Figure. 2J.
