## Supplementary Figure 6 for "Human pluripotent stem cell-derived intestinal epithelial cells maintain small intestine-specific functions over time, even with repeated cell division"

Phase I

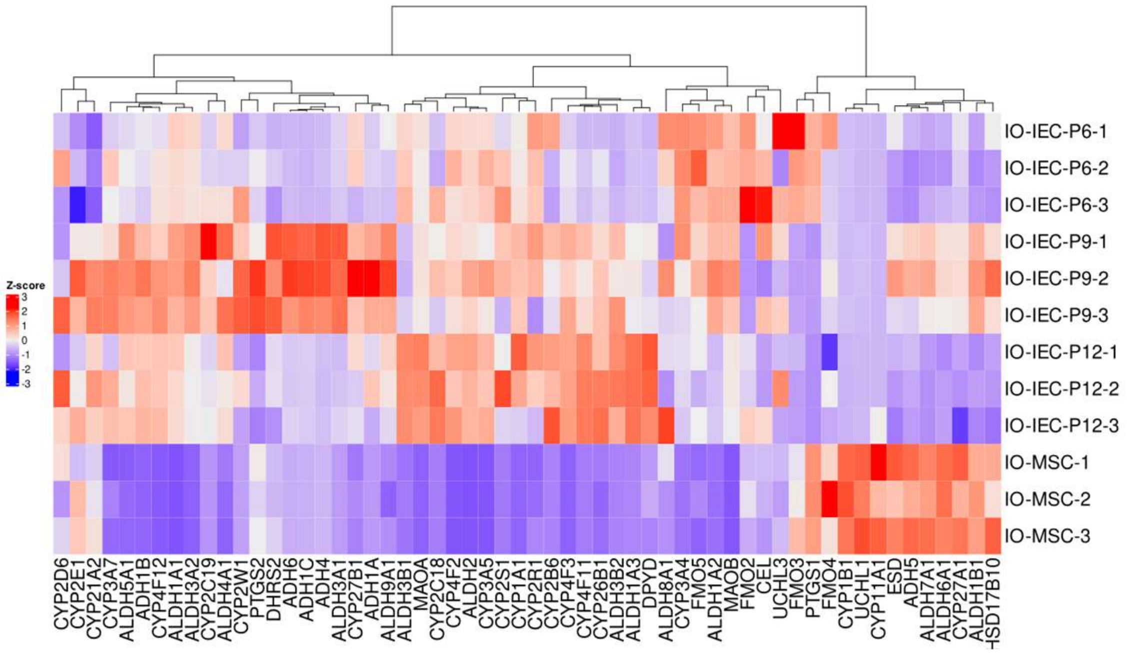


Phase II

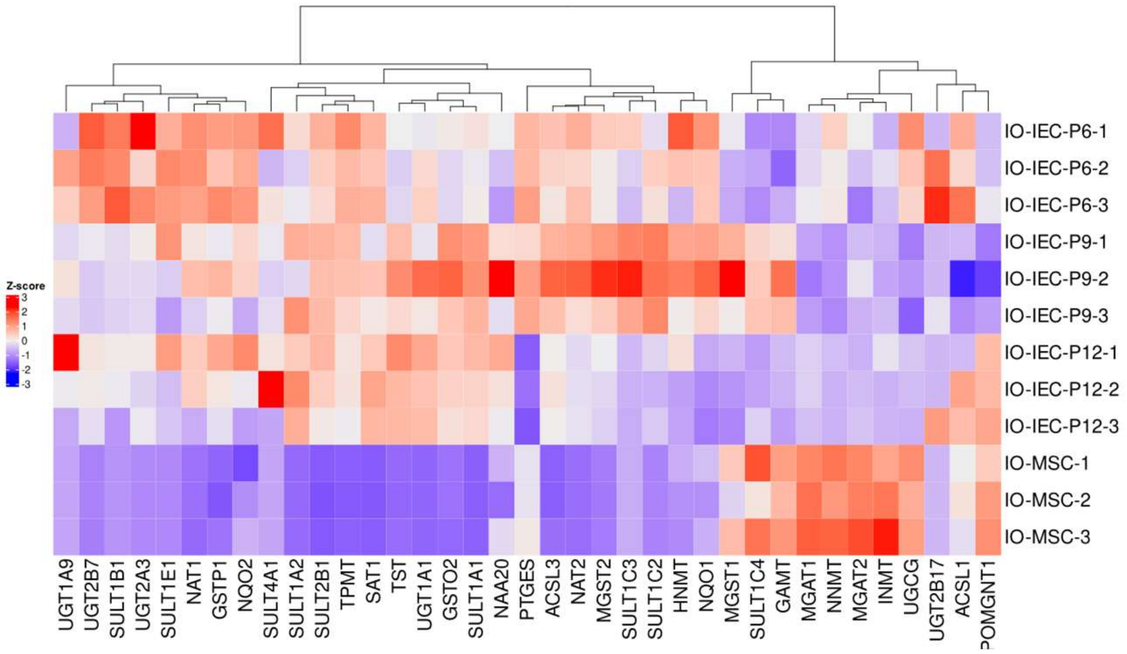


**Supplementary Figure 6. Drug metabolism-related genes in organoid-derived intestinal epithelial cells at each passage**

**Supplementary Figure 6. Drug metabolism-related genes in organoid-derived intestinal epithelial cells at each passage**

Intestinal organoid-derived intestinal epithelial cells are abbreviated as IO-IEC. Intestinal organoid-derived mesenchymal stromal cells are abbreviated as IO-MSC. Overview of drug metabolism-related gene expression in intestinal organoid-derived intestinal epithelial cells.
