## Supplementary Figure 8 for "Human pluripotent stem cell-derived intestinal epithelial cells maintain small intestine-specific functions over time, even with repeated cell division"

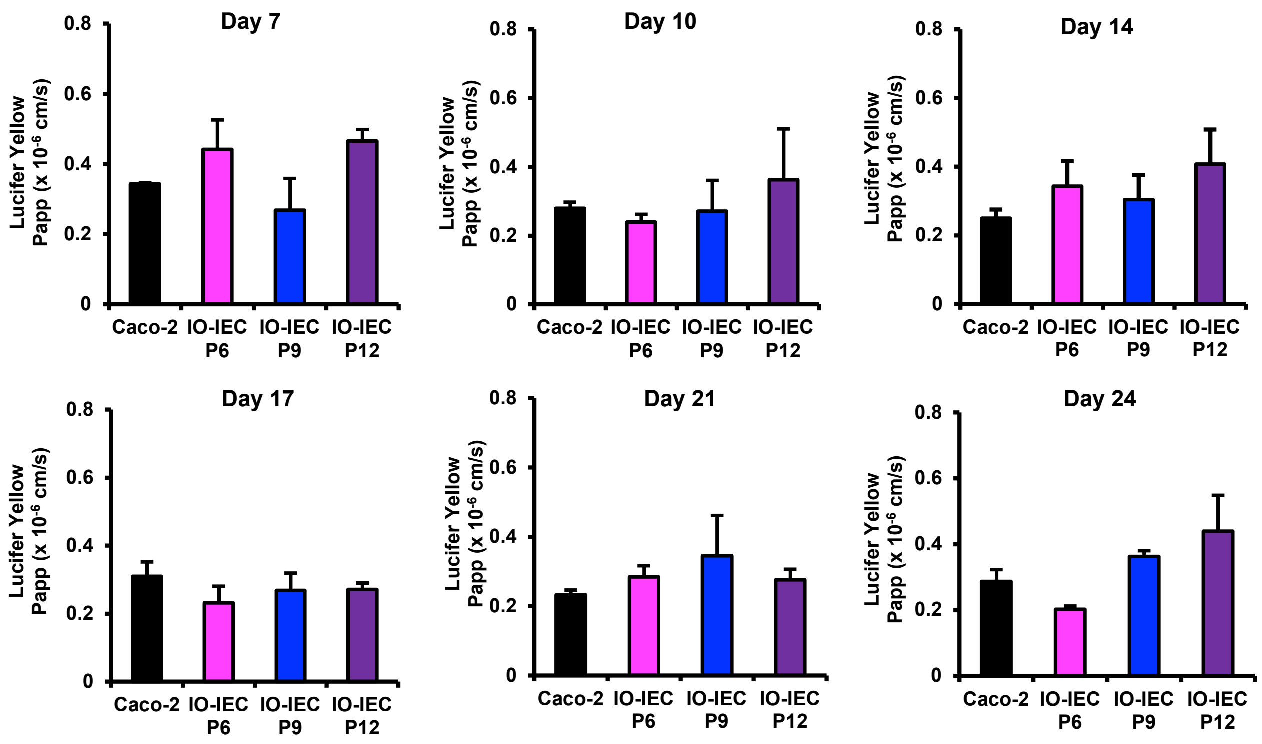


**Supplementary Figure 8. Lucifer Yellow values over 4 weeks cultured for organoid-derived intestinal epithelial cells**

**Supplementary Figure 8. Lucifer Yellow values over 4 weeks cultured for organoid-derived intestinal epithelial cells**

Intestinal organoid-derived intestinal epithelial cells are abbreviated as IO-IEC. Intestinal organoid-derived mesenchymal stromal cells are abbreviated as IO-MSC. Lucifer Yellow permeability test of organoid-derived intestinal epithelial cells at Passage 6, 9, and 12, intestinal organoid-derived mesenchymal stromal cells, and Caco-2 cells each day. Results are expressed as mean ± SD (n = 3 triplicate biological experiments).
