## Supplementary Figure 10 for "Human pluripotent stem cell-derived intestinal epithelial cells maintain small intestine-specific functions over time, even with repeated cell division"

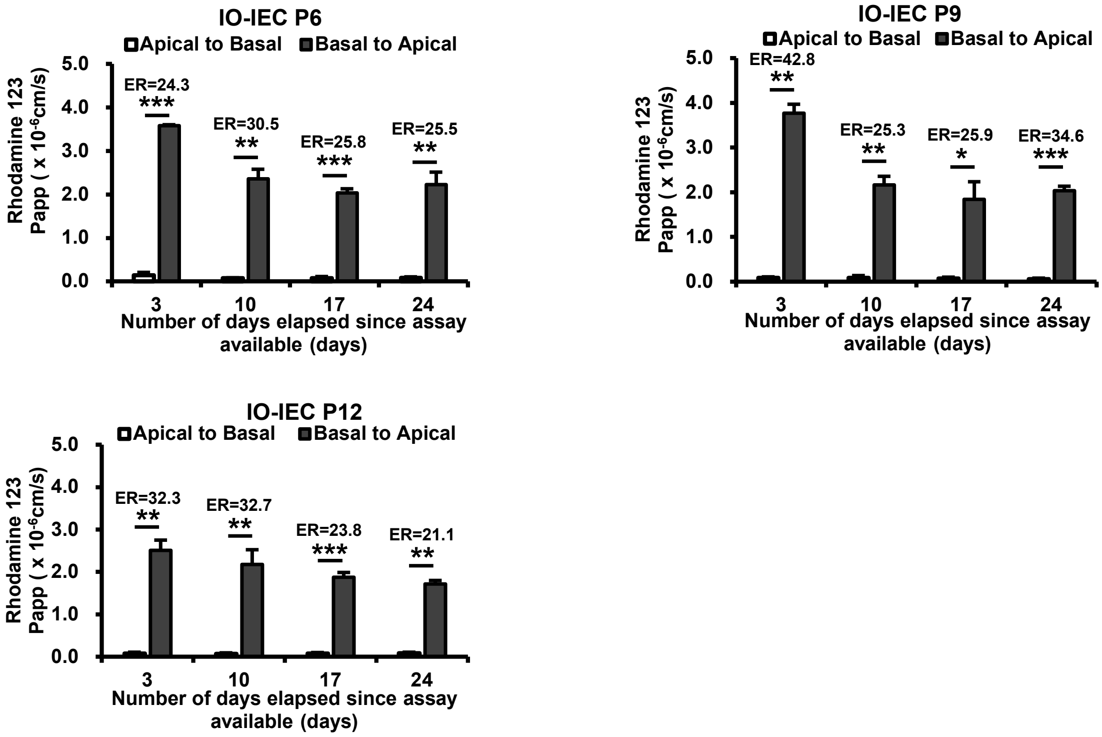


**Supplementary Figure 10. The p-gp activity over 4 weeks of culture for organoid-derived intestinal epithelial cells at each passage**

**Supplementary Figure 10. The p-gp activity over 4 weeks of culture for organoid-derived intestinal epithelial cells at each passage**

Intestinal organoid-derived intestinal epithelial cells are abbreviated as IO-IEC. Permeability test with Rhodamine 123. The efflux ratio (ER) for Rhodamine 123 was derived from the Papp values associated with basal-to-apical transport (B to A) and apical-to-basal transport (A to B). organoid-derived intestinal epithelial cells at Passage 6, 9, and 12. Results are expressed as mean ± SD (n = 3 triplicate biological experiments). Statistical significance was determined using Student's t-test; *** $p < 0.001$, ** $p < 0.01$, * $p < 0.05$.
