## Supplementary Figure 13 for "Human pluripotent stem cell-derived intestinal epithelial cells maintain small intestine-specific functions over time, even with repeated cell division"

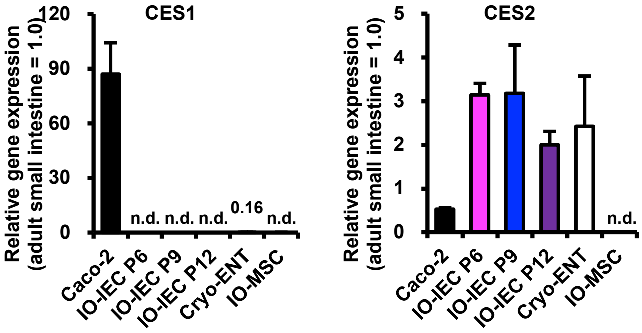


**Supplementary Figure 13. Carboxylesterase expression at 4 weeks**

**Supplementary Figure 13. Carboxylesterase expression at 4 weeks**

Intestinal organoid-derived intestinal epithelial cells are abbreviated as IO-IEC. Intestinal organoid-derived mesenchymal stromal cells are abbreviated as IO-MSC. Cryopreserved human enterocytes are abbreviated as Cryo-ENT. CES1 and CES2 gene expressions by qRT-PCR analysis in organoid-derived intestinal epithelial cells, intestinal organoid-derived mesenchymal stromal cells, and Caco-2 cells at 4 weeks and cryopreserved human enterocytes. The expression level of adult small intestine whole tissue was set at 1.0. Results are expressed as mean ± SD (n = 3 triplicate biological experiments).
