## Supplementary Figure 14 for "Human pluripotent stem cell-derived intestinal epithelial cells maintain small intestine-specific functions over time, even with repeated cell division"

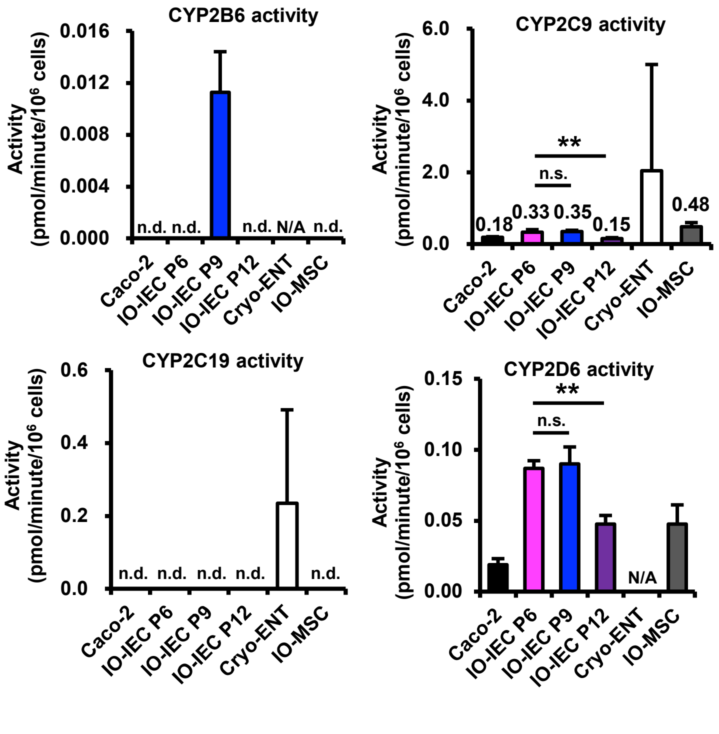


**Supplementary Figure 14. Cytochrome P450 activity in organoid-derived intestinal epithelial cells**

**Supplementary Figure 14. Cytochrome P450 activity in organoid-derived intestinal epithelial cells**

Intestinal organoid-derived intestinal epithelial cells are abbreviated as IO-IEC. Intestinal organoid-derived mesenchymal stromal cells are abbreviated as IO-MSC. Cryopreserved human enterocytes are abbreviated as Cryo-ENT. Measure Cytochrome P450 (CYP2B6, 2C9, 2C19, and 2D6) activity in Caco-2 cells, organoid-derived intestinal epithelial cells, and intestinal organoid-derived mesenchymal stromal cells. Results are expressed as mean ± SD (n = 3 triplicate biological experiments, cryopreserved human enterocytes (n = 6). Statistical significance was determined using one-way ANOVA with Dunnett's test compared to organoid-derived intestinal epithelial cells Passage 6. *** $p < 0.001$, ** $p < 0.01$, * $p < 0.05$. n.s.: not significant. n.d.: not detected. N/A: no data.
