## Supplementary Figure 15 for "Human pluripotent stem cell-derived intestinal epithelial cells maintain small intestine-specific functions over time, even with repeated cell division"

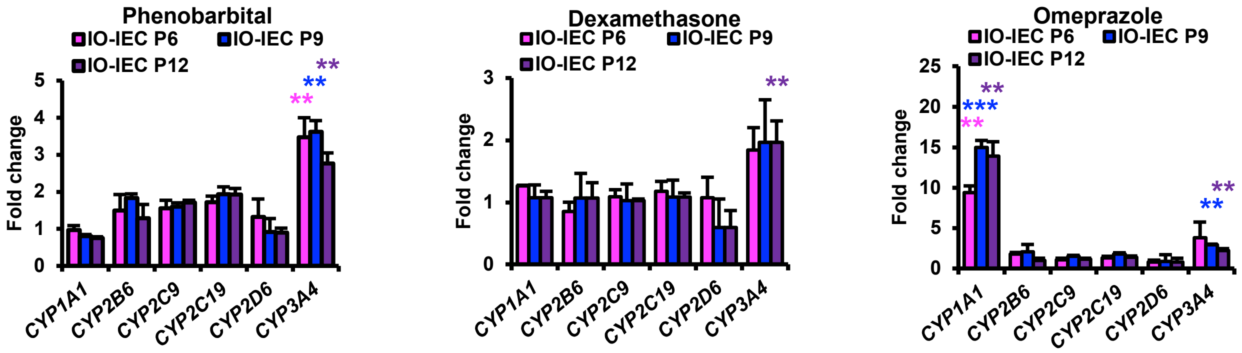


**Supplementary Figure 15. Induction of the cytochrome P450 genes in organoid-derived intestinal epithelial cells**

**Supplementary Figure 15. Induction of the cytochrome P450 genes in organoid-derived intestinal epithelial cells**

Intestinal organoid-derived intestinal epithelial cells are abbreviated as IO-IEC. Induction of the cytochrome P450 genes with exposure to phenobarbital, dexamethasone, and omeprazole. Results are expressed as mean ± SD (n = 3). The expression level of each gene without any treatment (DMSO) was set to 1.0. Expression levels were calculated from the results of independent (biological) triplicate experiments. Statistical significance was determined using Student's t-test; *** Fold > 2, $p < 0.001$, ** Fold > 2, $p < 0.01$, * Fold > 2, $p < 0.05$.
